# Long-read-sequenced reference genomes of the seven major lineages of enterotoxigenic *Escherichia coli* (ETEC) circulating in modern time

**DOI:** 10.1101/2020.07.16.203430

**Authors:** Astrid von Mentzer, Grace A. Blackwell, Derek Pickard, Christine J. Boinett, Enrique Joffré, Andrew J Page, Ann-Mari Svennerholm, Gordon Dougan, Åsa Sjöling

## Abstract

**Background:** Enterotoxigenic *Escherichia coli* (ETEC) is an enteric pathogen responsible for the majority of diarrheal cases worldwide. ETEC infections are estimated to cause 80,000 fatalities per year, with the highest rates of burden, ca 75 million cases per year, amongst children under five years of age in resource-poor countries. It is also the leading cause of diarrhoea in travellers. Previous large-scale sequencing studies have found seven major ETEC lineages currently in circulation worldwide.

**Results:** We used PacBio long-read sequencing combined with Illumina sequencing to create high-quality complete reference genomes for each of the major lineages with manually curated chromosomes and plasmids. The plasmids carrying ETEC virulence genes were compared to other available long-read sequenced ETEC strains using blastn. The ETEC reference strains harbour between two and five plasmids, including virulence, antibiotic resistance and phage-plasmids. The virulence plasmids carrying the colonisation factors are highly conserved as shown by comparison with plasmids with other ETEC strains and confirm that the plasmids and chromosomes of ETEC are both crucial for ETEC virulence and success as pathogens.

**Conclusion:** We confirm that the major ETEC lineages all harbour conserved plasmids that have been associated with their respective background genomes for decades. The in-depth analysis of gene content, synteny and correct annotations of plasmids will elucidate other plasmids with and without virulence factors in related bacterial species. These reference genomes allow for fast and accurate comparison between different ETEC strains, and these data will form the foundation of ETEC genomics research for years to come.

## Background

Diarrheal pathogens are a leading cause of morbidity and mortality globally (WHO, 2017), with enterotoxigenic *Escherichia coli* (ETEC) accounting for a large proportion of the diarrhoea cases in resource-poor countries [1]. An estimation of 220 million cases each year are attributed to ETEC (WHO PPC 2020). The most vulnerable group is children under five years, but ETEC can also cause disease in adults and is the principal cause of diarrhoea in travellers. Resource-poor settings, where access to clean water is limited, enable the spread of ETEC, transmitted via the faecal-oral route through ingestion of contaminated food or water [2]. The disease severity may range from mild to cholera-like symptoms with profuse watery diarrhoea. The infection is usually self-limiting, lasting three to four days and may be treated by water and electrolyte rehydration to balance the loss of fluids and ions. There is strong evidence to support that an ETEC vaccine is of key importance to prevent children and adults from developing ETEC disease [3]. Several efforts are on-going to develop an ETEC vaccine, with the majority focusing on including immunogenic antigens possibly capable of inducing protection against a majority of the circulating ETEC clones [4–6].

ETEC bacteria adhere to the small intestine through fimbrial, fibrillar or afimbrial outer membrane-structures called colonisation factors (CF). Upon colonisation, the bacteria proliferate and secrete heat-labile toxin (LT) and/or heat-stable toxins, (STh or STp) causing diarrhoea and often vomiting causing the further spread of the bacteria in the environment [7].

The ability of an ETEC strain to infect relies on its ability to adhere to cells of a specific host. To this day, 27 different CFs with human tropism have been described, and individual ETEC strains usually express 1-3 different CFs [8–14]. The enterotoxins, LT and ST, can also be subdivided based on structure and function. Human-associated ETEC strains express one of the 28 different LT-I variants (LTh-1 and LTh-2 are the most common variants) [15] alone or together with one of the genetically distinct types of STa; STh and STp [16,17].

We have previously shown that ETEC strains causing human disease can be grouped into a set of clonal lineages that encompass strains with specific virulence profiles. Seven of the 21 identified lineages encompass ETEC strains that express the most commonly found CFs and toxin profiles amongst isolated clinical ETEC strains [4,18].

There is currently one complete ETEC reference genome, H10407 [19], with curated annotations. Several additional complete ETEC genomes are available [20,21], some of which are annotated using automated annotation pipelines that often fail at correctly annotating ETEC specific genes such as CFs. The rapid adaptation of next-generation sequencing in public health, specifically within bacterial diseases [22,23] and several large-scale sequencing studies [18,24–28] has led to a sharp increase in the number of publicly available ETEC genomes. Most of these data were generated with short-read technologies, such as Illumina. A limitation of short-read sequence data is the inability to unambiguously resolve repetitive regions of a genome, leading to fragmented de novo assemblies of the underlying genome, missing regions and genes, and disjointed synteny. ETEC is a highly diverse pathogen both in the core genome (SNPs) and the accessory genome, including mobile genetic elements (MGE). Clinically related MGEs, such as virulence plasmids, vary within ETEC strains. Hence, it is important to identify lineage-specific reference genomes that are carefully annotated, i.e. manually curated annotations, and include both chromosome and plasmid(s). Several complete genomes have been generated using long-read sequencing alone [20,28], however, circularising some chromosomes and plasmids may be difficult, and small plasmids can be lost. Assembly issues can be resolved using a hybrid assembly approach combining long-read and short-read sequencing data. In this report, we describe eight genomes, eight chromosomes (seven successfully circularised) and 29 plasmids (24 successfully circularised) with curated annotations, from isolates representing the major ETEC lineages (L1-L7) that cause disease globally. They are sequenced using both short and long-read sequencing technologies to provide the highest accuracy currently available. These reference genomes will form the foundation of ETEC genomics research for years to come.

## Results

### Genome analysis of eight representative ETEC isolates

Eight ETEC strains representing the seven major ETEC lineages (L1-L7) comprising isolates with the most prevalent virulence factor profiles were sequenced, assembled, circularised and manually curated (Table 1).

**Table 1:** Characteristics of the reference ETEC strains.

| Strain | Lineage | Phylo-group | MLST | O antigen | CF | Toxin profile | YoI<sup>a</sup> | Location | Subject | Age of subject | D/AS<sup>b</sup> |
| --- | --- | --- | --- | --- | --- | --- | --- | --- | --- | --- | --- |
| E925 | L1 | A | 2353 | O6 | CS1+CS3+CS21 | LT+STh | 2003 | Guatemala | Indigenous | Child <5 yrs | D |
| E1649 | L2 | A | 4 | O6 | CS2+CS3+CS21 | LT+STh | 1997 | Indonesia | Traveller | Adult | D |
| E36 | L3 | B1 | 173 | O78 | CFA/I+CS21 | LT+STh | 1980 | Bangladesh | Indigenous | Adult/child | D |
| E2980 | L3 | B1 | 5305 | O114 | CS7 | LT | 2010 | Bangladesh | Indigenous | Child <5 yrs | D |
| E1441 | L4 | A | 1312 | O25 | CS6+CS21 | LT | 1997 | Kenya | Traveller | Adult | D |
| E1779 | L5 | B1 | 443 | O115 | CS5+CS6 | LT+STh | 2005 | Bangladesh | Indigenous | Adult | D |
| E562 | L6 | A | 2332 | ON3 | CFA/I+CS21 | STh | 2000 | Mexico | Traveller | Adult | D/AS |
| E1373 | L7 | E | 182 | O169 | CS6 | STp | 1996 | Indonesia | Traveller | Adult | D |

<sup>a</sup> YoI: Year of isolation
<sup>b</sup> D/AS: Diarrhoea or Asymptomatic

L3 includes two different representative strains, one CS7 and one CFA/I positive strain. All chromosomes except one were circularised (E1779). The average length of the chromosome was 4,927,521 bases (4,721,269-5,151,162) with an average GC content of 50.7% (50.4% to 50.9%) and the number of CDS ranging from 4,409 to 4,924 (Table S1). Each ETEC reference genome contains between two and five plasmids encompassing plasmid-specific features. Some of which carried virulence genes and/or antibiotic resistance genes (Table 2, Additional File 2).

**Table 2:**
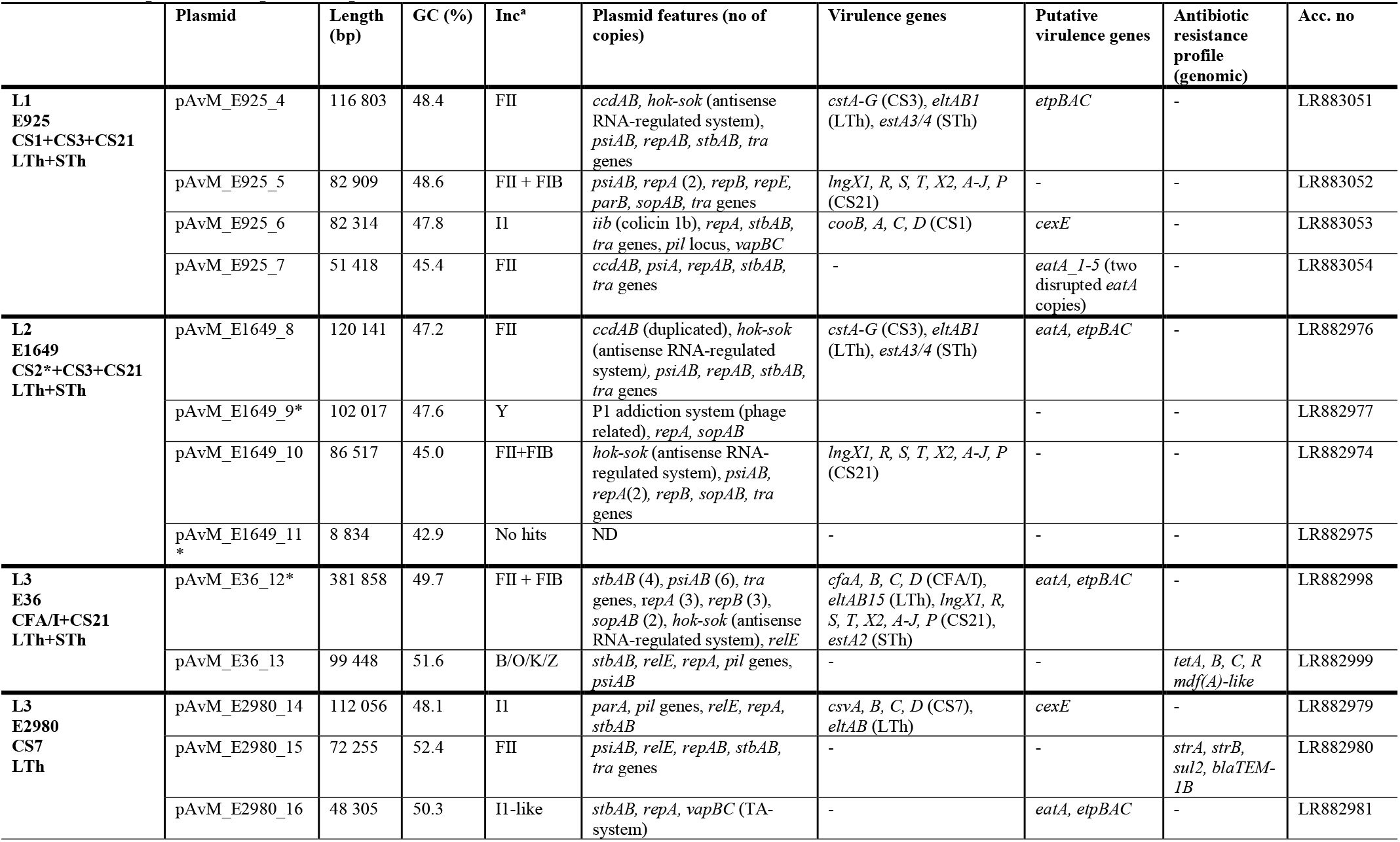
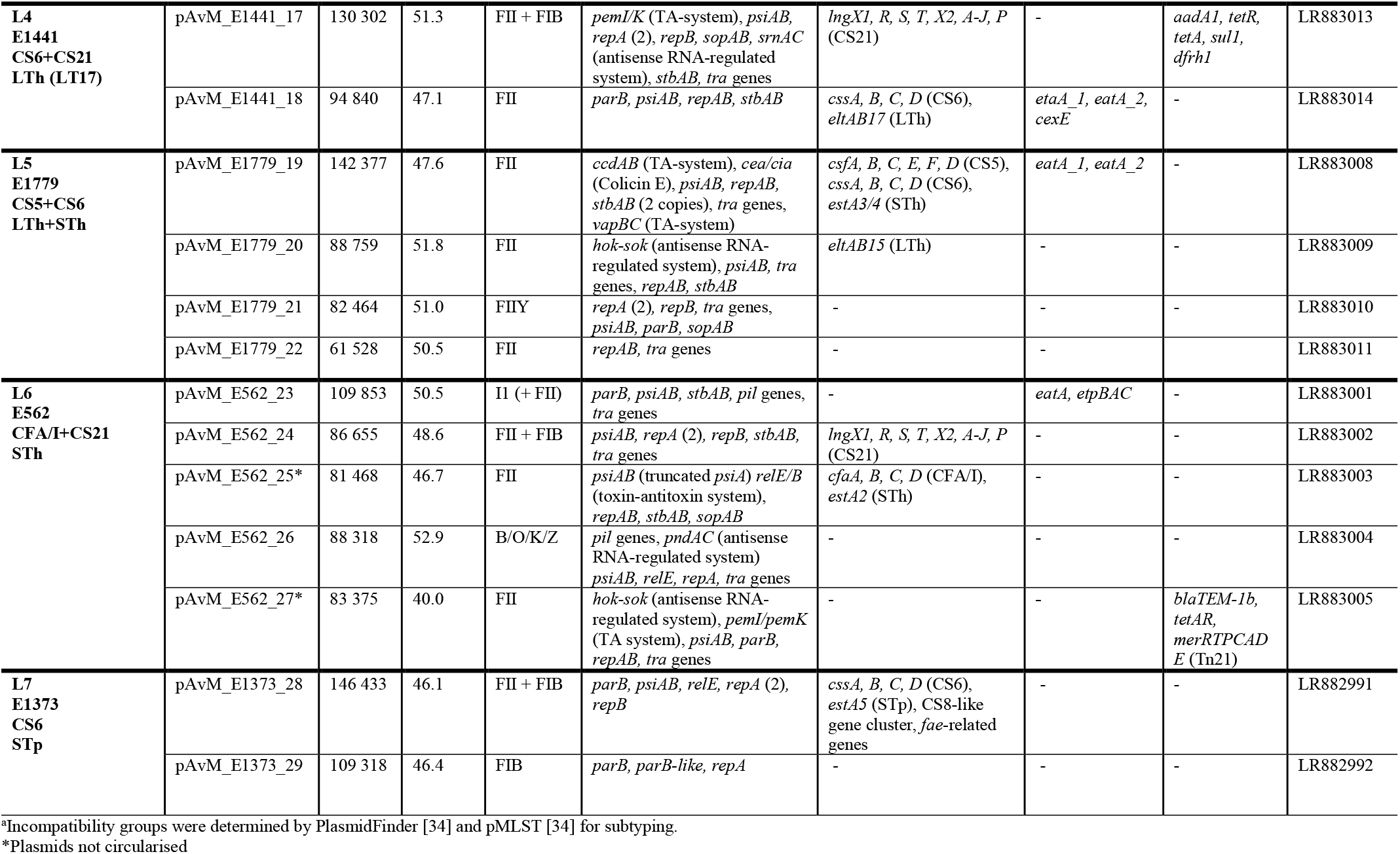
Description of the plasmids present in the 8 ETEC reference strains.

### Comparative genomics of the chromosome

The chromosomes of the reference strains were aligned and compared using MAUVE, and the overall structure is conserved across all eight chromosomes (Figure S1). In total, 8,348 chromosomal genes were identified in the eight ETEC strains with 3,179 genes considered part of the core genome shared by all eight reference strains. The majority of human commensal *Escherichia coli* (*E. coli*) strains belong to subgroup A [29,30]. However, ETEC strains fall into multiple phylogenetic groups (A, B1, B2, D, E, F and CladeI with the majority found in the phylogenetic groups A and B1 [18]. The phylogenetic group of the eight ETEC reference strains have previously been determined using the triplex-PCR scheme [31]. The ETEC references were re-analysed using ClermonTyping [32] and it was determined that strain E1373 belongs to the phylogenetic group E while the other reference isolates belong to groups A and B1 (Table 1).

### Plasmids

The plasmids of each isolate were annotated using Prokka followed by manual curation of the annotations including genes part of the conjugation machinery and known plasmid stability genes. Virulence factors (including CFs, toxins and putative virulence factors), antibiotic resistance determinants with the Comprehensive Antibiotic Resistance Database (CARD) [33] as well as complete and partial insertion elements and prophages were manually annotated. The plasmids were designated pAvM_strainID_integer, e.g. pAvM_E925_4 (Additional file 2). The first plasmid reported in this study starts at 4 as three previous plasmids E873p1-3 already have been deposited to GenBank related to a different project [8].

Plasmids were typed by analysing the presence and variation of specific replication genes to assign the plasmids to incompatibility (Inc) groups. The Inc groups of the ETEC reference plasmids were first determined using PlasmidFinder and further classified into subtypes using pMLST [34]. The replicons identified are IncFII, IncFIIA, IncFIIS, IncFIB, IncFIC, IncI1 and IncY. Plasmids with replicon IncY, IncFIIY or IncB/O/K/Z mainly harboured plasmid associated genes, such as stability and transfer genes. Importantly, replicons FII, FIB and I1 were found to be strongly associated with virulence genes as all CF loci, toxins and virulence factors *eatA* and *etpBAC* were present on these plasmids. The majority of all ETEC plasmids analysed here (17/29) belong to IncFII, of which six of the IncFII plasmids have an additional IncFIB replicon. In six of the ETEC reference strains two or three IncFII replicons are present, for example, in strain E925, the plasmids pAvM_E925_4 and 7 both belong to IncFII. However, the plasmids were further subtyped to FII-111 and FII-15, respectively, (Table 2 and Additional file 3), explaining the plasmid compatibility.

### Virulence factors

The CFs expressed by the selected reference strains are CFA/I, CS1-CS3, CS5-CS7 and CS21. Three of the strains (E925, E1649 and E1779) express both LT and ST, two strains (E2980 and E1441) express LT and the strains E36 and E562 express STh, while E1373 express STp (Table 1). A plasmid can harbour multiple virulence genes, usually a CF locus and genes encoding one or two toxins. Interestingly, plasmids do not often harbour multiple CF loci, but on individual plasmids (in the ETEC reference strains described here). Exceptions for this is strain E1779 in which CS5 and CS6 loci are located on the same plasmid (pAvM_E1779_19). In both E925 (L1) and E1649 (L2) the genes encoding CS3 (*cstABGH*), ST (*estA*) and LT (*eltAB*) are located on the same plasmid, both with the FII replicon and of roughly the same size (Table 2). Blastn comparison between the plasmids and additional plasmids that harbour the same virulence genes shows that they are highly conserved (Figure 1). The results correspond with the close genetic relationship and common ancestry of lineage 1 (L1) and lineage 2 (L2) [18].

**Figure 1:**
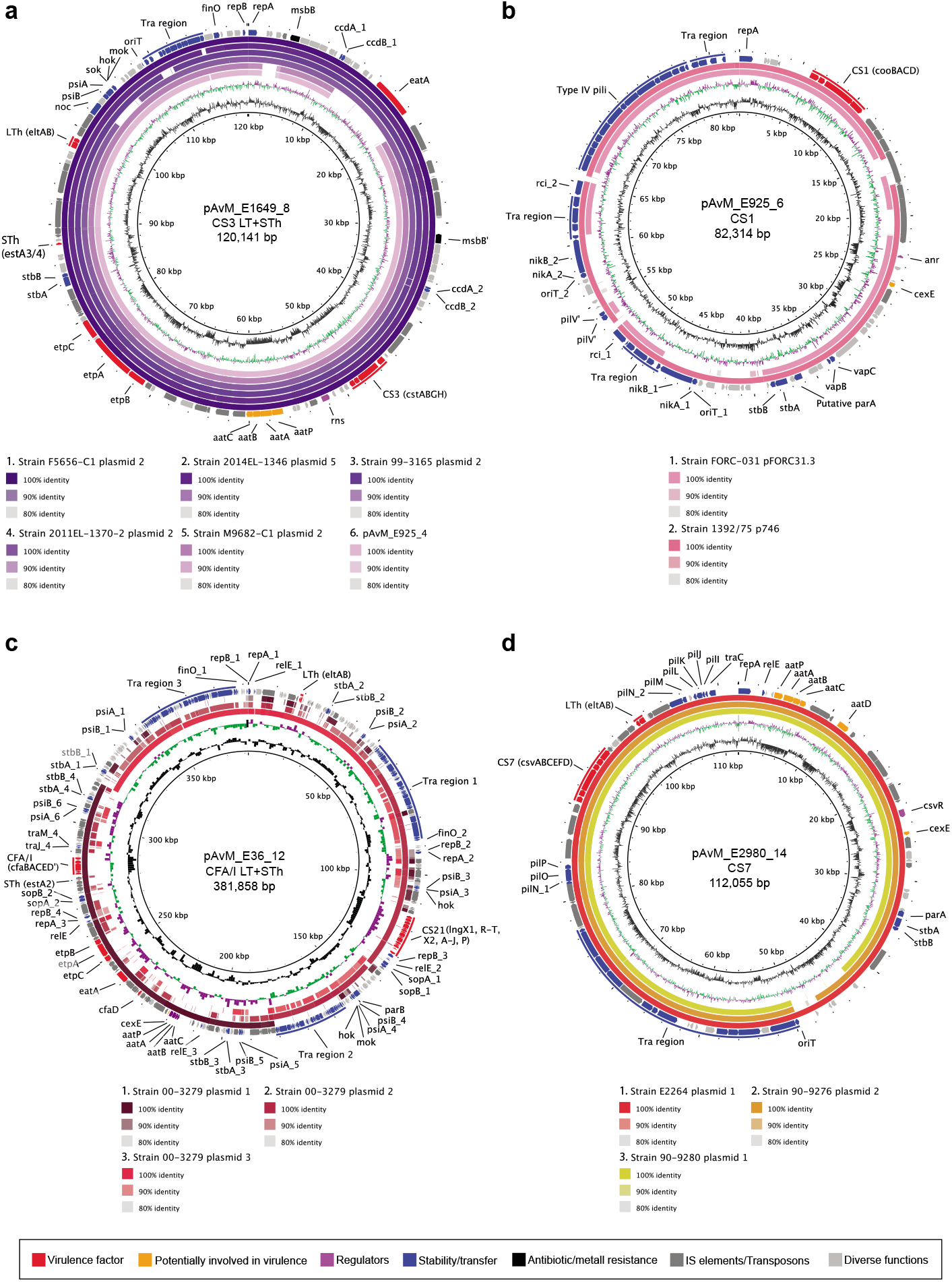
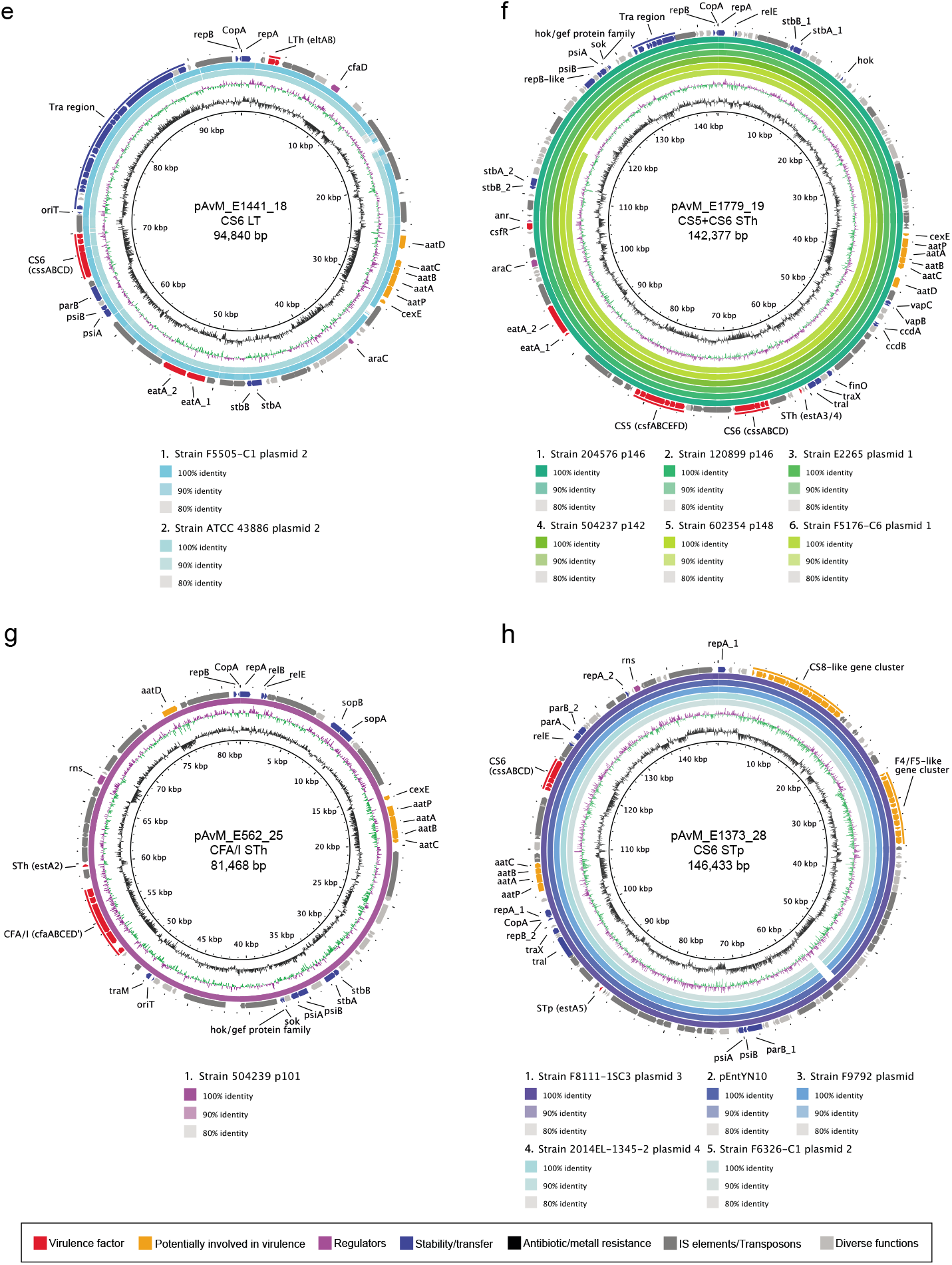
Comparison between the ETEC reference plasmids harbouring colonisation factors and other PacBio-sequenced ETEC plasmids using blastn. **a)** pAvM_E1649_8 (CS3) as reference and pAvM_925_4 (CS3) compared to the following ETEC plasmids: F5656-C1 plasmid 2 (USA, CP024262.1), 2014-EL-1346-6 plasmid 5 (2014, USA; CP024237.1), 99-3165 plasmid 2 (USA; CP029980.1), 2011EL-1370-2 plasmid 2 (2011, USA; CP022914.1) and M9682-C1 plasmid 2 (1975, USA; CP024277.1). **b)** pAvM_E925_6 (CS1) compared to ETEC plasmids pFORC31.3 (2004, Korea; CP013193.1) and 1392/75 p746 (1973; FN822748.1). **c)** pAvM_E36_12 (CFA/I) compared to plasmids 1-3 (p1: CP024294.1; p2 CP024295.1; p3: CP024296.1) from ETEC strain 00-3279 (USA). **d)** pAvM_E2980_14 (CS7) compared to E2264 plasmid 1 (2006, Bangladesh; CP023350.1), 90-9276 plasmid 2 (1988, Bangladesh; CP024298.1) and 90-9280 plasmid 1 (1988, Bangladesh; CP024241.1). **e)** pAvM_E1441_18 (CS6) compared to F5505-C1 plasmid 2 (2013, Sweden; CP023259.1) and ATCC 43886 plasmid 2 (CP024255.1). **f)** pAvM_E1779_19 (CS5+CS6) compared to 204576 p146 (2010, Mali; CP025908.1), 120899 p146 (2012, Gambia; CP025917.1), E2265 plasmid 1 (2006, Bangladesh; CP023347.1), 504237 p142 (2010, India; CP025863.1), 602354 p148 (2009, Bangladesh; CP025848.1) and F5176-C6 plasmid 1 (1997; CP024668.1). **g)** pAvM_E562_25 (CFA/I) compared to p504239_101 (2010, India; CP025860.1). **h)** pAvM_E1373_28 (CS6) compared to F8111-1SC3 plasmid 3 (USA; CP024272.1), pEntYN10 (1991, Japan; AP014654.2), F9792 plasmid (USA; CP023274.1), 2014EL-1345-2 plasmid 4 (2014, USA; CP024227.1) and F6326-C1 plasmid 2 (1998, USA; CP024265.1). The thresholds chosen for the blastn are shown in the key below each plasmid comparison. The colour code for the annotations are listed at the bottom of the figure. The two most inner rings depict GC content in black and GC Skew-in purple and GC Skew+ in green. The figures were generated using BRIG [84] v0.95.

Besides CFs and toxins, additional virulence factors were identified in the majority of the strains (Table 2), with *eatA* and *etpBAC* being the most commonly found.

EatA is an immunogenic mucinase that contributes to virulence by degrading MUC2 which is the major protein component of mucus in the small intestine [35,36]. The *etpABC* genes encode an adhesin located on the tip of the flagella and mediate adherence to host cells [37,38]. Four reference strains (E925, E1649, E36 and E562) harbour both *eatA* and *etpBAC*. In three strains the *eatA* and/or *etpBAC* are located on the same plasmid with an FII or FII+FIB replicon along with additional ETEC virulence genes, except in E562 and E1373, where *eatA* and *etpBAC* are located on an I1+FII (pAvM_E562_23) and I1 (pAvM_E1373_16) plasmid, respectively, which mainly contains plasmid associated genes including genes encoding the *pil* operon and *tra*-operon (pAvM_E562_23). Furthermore, a less explored putative virulence factor is CexE, which is an extracytoplasmic protein dependent on the expression of the CFA/I regulator *cfaD* [39], and was first identified in H10407 [40]. Corroborating earlier findings, the CFA/I positive E36 (L3) and E562 (L6) isolates harbour *cexE* (pAvM_E36_12 and pAvM_E562_25). In addition, *cexE* is present in pAvM_E925_6, pAvM_E1779_19 and pAVM_E2980_14, pAvM_E1441_18 and pAvM_E1373_28. *CexE* has previously also been identified in several CS5+CS6 positive ETEC and shown to be upregulated in the presence of bile and sodium glycocholate-hydrate [41]. Bile is known to be involved in the regulation of several ETEC CFs [42,43]. The location of *cexE* seems to be conserved across specific strains. In pAvM_E36_12, pAvM_E1441_18, pAvM_E1779_19 and pAvM_E562_25 *cexE* is located upstream of the *aatPABC* locus, whereas in pAvM_E925_6 and pAvM_E2980_14 *cexE* is located downstream of *rob* (an AraC family transcriptional regulator) in the opposite direction. pAvM_E925_4 harbours the *aatPABC* locus; however *cexE* is located on a different plasmid (pAvM_E925_6) in this strain.

### Comparison of plasmids with the same virulence profile

ETEC isolates within a lineage share the same virulence profile, specifically the same CF profile (Figure S10). We verified that our selected isolates grouped within previously described lineages with confirmed virulence profiles by phylogenetic analyses (Figure S2). Blastn of each of the CF positive plasmids from each reference genome were performed, and the best hit(s) were used for subsequent analysis (Figure 1). Most of the plasmids identified as related to the ETEC reference plasmids were not annotated, hence, when needed these were annotated using the corresponding ETEC reference plasmids annotation as a high priority when running Prokka. We show that plasmids with the same CF and toxin profile from the same lineage are often conserved (Figure 1). For example, the two plasmids encoding CS3 (pAvM_E925_4 and pAvM_E1649_8) are highly similar to several CS3 harbouring plasmids from O6:H16 strains collected from various geographical locations between 1975 and 2014, including *E. coli* O6:H16 strain M9682-C1 plasmid unnamed2 (CP024277.1) and *E. coli* strain O6:H16 F5656C1 plasmid unnamed2 (CP024262.1) PacBio sequenced by Smith et al. [20] (Figure 1a). Furthermore, high coverage and similarity were found between the plasmids of isolates E1441 (L4), and PacBio sequenced plasmids of ETEC isolates ATCC 43886/E2539C1 and 2014EL-1346-6 [20]. These isolates were collected in the seventies [44] and 2014 (from a CDC collection), respectively, and assigned as O25: H16 which is the O group determined for E1441 in silico (Figure 1e). Plasmids of E2980 (LT+ CS7, L3) were validated by the PacBio sequenced plasmids of ETEC isolate E2264 (Figure 1d). Similarly, two plasmids of E1779 (LT, STh+CS5+CS6, L5) was identified in E2265 (LT, STh+CS5+CS6 [28,41], although E1779 harboured two additional plasmids. Several additional L5 ETEC genomes have been sequenced within the GEMS study [45], and high plasmid similarity and conservation in CS5+CS6 positive L5 isolates was evident (Figure 1f).

Overall the results show that ETEC plasmids are specific to lineages circulating worldwide and conserved over time (Figure 1, Figure S2-S9 for more extensive plasmid annotation and Figure S10). Thus, the plasmids of major ETEC lineages must confer evolutionary advantages to their host genomes since they are seldom lost.

### Antibiotic resistance

*E. coli* can become resistant to antibiotics, both via the presence of antibiotic resistance genes and the acquisition of adaptive and mutational changes in genes encoding efflux pumps and porins which allows the bacterium to pump out the antibiotic molecules effectively.

Antibiotic resistance genomic marker(s), both chromosomally located and on plasmids, were identified using the CARD database [33] (Table 2, Figure S12 and Figure S13 and Additional file 2). Similar to other studies, IncFII and B/O/K/Z plasmids were found to harbour genes conferring antibiotic resistance [46]. Furthermore, the phenotypic antibiotic resistance profile was determined with clinical MIC breakpoints based on EUCAST (The European Committee on Antimicrobial Susceptibility Testing) [47] (Table S2). Phenotypic antibiotic resistance profiles (Table S2) were supported mainly by the findings of antibiotic resistance genes, efflux pumps and porins (Figures S4 and S5 and Table S3), although some differences were found. All ETEC reference strains are phenotypically resistant to at least two antibiotics of the 14 tested (Table S2). Resistance against penicillin’s, norfloxacin (Nor) and chloramphenicol (Cm) is most common among these strains. Two of the strains, E1441 and E2980, harbour more than four antibiotic resistance genes as well as multiple efflux systems and porins (Figure S12, Figure S13 and Table S3). The plasmid pAvM_E1441_17 carries *aadA1-like, dfrA15, sul1* and *tetA*(A) resistance genes (Table 2), where the first three genes are in a Class 1 integron which confers resistance to streptomycin, trimethoprim, and sulphonamide (sulphamethoxazole). The gene *tetA*(A) is part of a truncated Tn*1721* transposon [48]. The E1441 strain was verified as resistant to tetracycline (Tet) and sulphamethoxazole-trimethoprim (Sxt) while streptomycin was not tested. A *mer* operon derived from Tn*21* is also present the resistance region of pAvM_E1441_17 (Table 2), indicating that the plasmid would also likely confer tolerance to mercury, although this was not confirmed. Interestingly, this multi-replicon (FII and FIB) plasmid also harbours the *lng* locus encoding CS21, one of the most prevalent ETEC CFs. In isolate E2980 virulence plasmid pAvM_E2980_15 harboured multiple resistance genes in the same region (*bla*_TEM-1b_, *strA*, *strB* and *sul2*) conferring resistance to ampicillin, streptomycin and sulphonamides. E2980 was found to be resistant to ampicillin (Amp) and oxacillin (Oxa), which can be broken down by the beta-lactamase Bla_TEM-1b_, (Table 2, Table S2 and Table S3). E562 harbours three antibiotic resistance genes, *ampC* located in the chromosome and the *tet*(A) and *bla*_TEM-1b_ genes on an FII plasmid (pAvM_E562_27). The *mer* operon derived from Tn*21* is also present in the region (Table 2 and Table S3). The phenotypic resistance profile of E562 matches the genomic profile with resistance to tetracycline (Tet), ampicillin (Amp), amoxicillin-clavulanic acid (Amc) and oxacillin (Oxa) (Table S2). The plasmid pAvM_E36_13 contains a complete copy of Tn*10*, which encodes the *tet*(B), tetracycline resistance module. Although the AvM_E1373_29 phage-like plasmid is cryptic, related plasmids such as the pHMC2-family of phage-like plasmids [49] (described below), can harbour resistance genes such as *bla*_CTX-M-14_ [50] and *bla*_CTX-M-15_ [51,52].

Phenotypic intermediate resistance to ampicillin was found in E36 and E1779 encoded by chromosomal *ampC*. Higher MIC values against ampicillin are found in E2980 and E562 strains carrying *bla*_TEM_ genes. Phenotypic resistance to ceftazidime (Caz) and ceftriaxone (Cro) was not found in the isolates, which were consistent with the absence of extended-spectrum beta-lactamase (ESBL) resistance genes in the sequence data.

Resistance to chloramphenicol (Cm) was found in five isolates, but none of the resistant isolates contained known resistance genes suggesting that chromosomal mutations or presence of efflux pumps may account for this reduced susceptibility. The ETEC reference strains contain several efflux systems which could explain why the genotypic and phenotypic antibiotic resistance profile did not match for all antibiotics. All of the isolates harbour multiple efflux pumps located on the chromosome and plasmids (Table S3 and Figure S12). In E925, a non-synonymous mutation in *acrF* was identified (G1979A) resulting in a substitution from arginine to glutamine (A360Q). The effect on the expression and/or function of the AcrEF efflux pump was not verified.

Phenotypic resistance to norfloxacin (Nor) was found in 6 of the isolates. The isolates were analysed for chromosomal mutations likely to confer quinolone resistance, using ResFinder but mutations in *gyrA* were only found in one strain, E2980, at position S83A which may confer resistance to nalidixic acid, norfloxacin and ciprofloxacin. However, E2980 was sensitive to Nalidixic acid. Both mutation(s) that alter the target (*gyrA* and *parC*), as well as the presence of efflux pumps, can confer resistance to fluoroquinolones. The majority of the isolates are moderately resistant to Norfloxacin (and Nalidixic acid), both quinolones, which is most likely due to the presence of two efflux pumps, AcrAB-R and AcrEF-R, as only one mutation was identified in *gyrA* of isolate E2980 where usually at least two or more mutations are needed to confer augmented resistance [53].

### Identification of phage-like plasmids in ETEC

Two of the ETEC reference strains (E1649 and E1373) harboured phage-like plasmids (pAvM_E1649_9 and pAvM_E1373_29) which encode for DNA metabolism, DNA biosynthesis as well as structural bacteriophage genes (capsid, tail etc.). Both pAvM_E1649_9 and pAvM_E1373_29 contain genes associated with plasmid replication, division and maintenance (i.e. *repA* and *parAB*). Phage-like plasmids are found in various bacterial species, such as *E. coli*, *Klebsiella pneumoniae*, *Yersinia pestis*, *Salmonella enterica* serovar Typhi, *Salmonella enterica* serovar Typhimurium, *Salmonella enterica* serovar Derby and *Acinetobacter baumanii* [54]. The plasmid pAvM_E1649_9 belong to the P1 phage-like plasmid family (Figure 2a and Figure S14a) while pAvM_E1373_29 belongs to the pHCM2-family (Figure 2b and Figure S14b) that can be traced back to a likely phage origin similar to the *Salmonella* phage, SSU5 [49]. Both phage-plasmids thus contain replication and/or partition genes of plasmid origin and a complete set of genes that are phage related in function and properties (Figure 2 and Figure S14). Significantly, phage-like plasmid pAvM_E1373_29 falls more within the *E. coli* lineage of pHCM2 phage-like plasmid rather than those found in *Salmonella* species. This indicates that phage-like plasmids have diversified within the bacterial species they were isolated.

**Figure 2:**
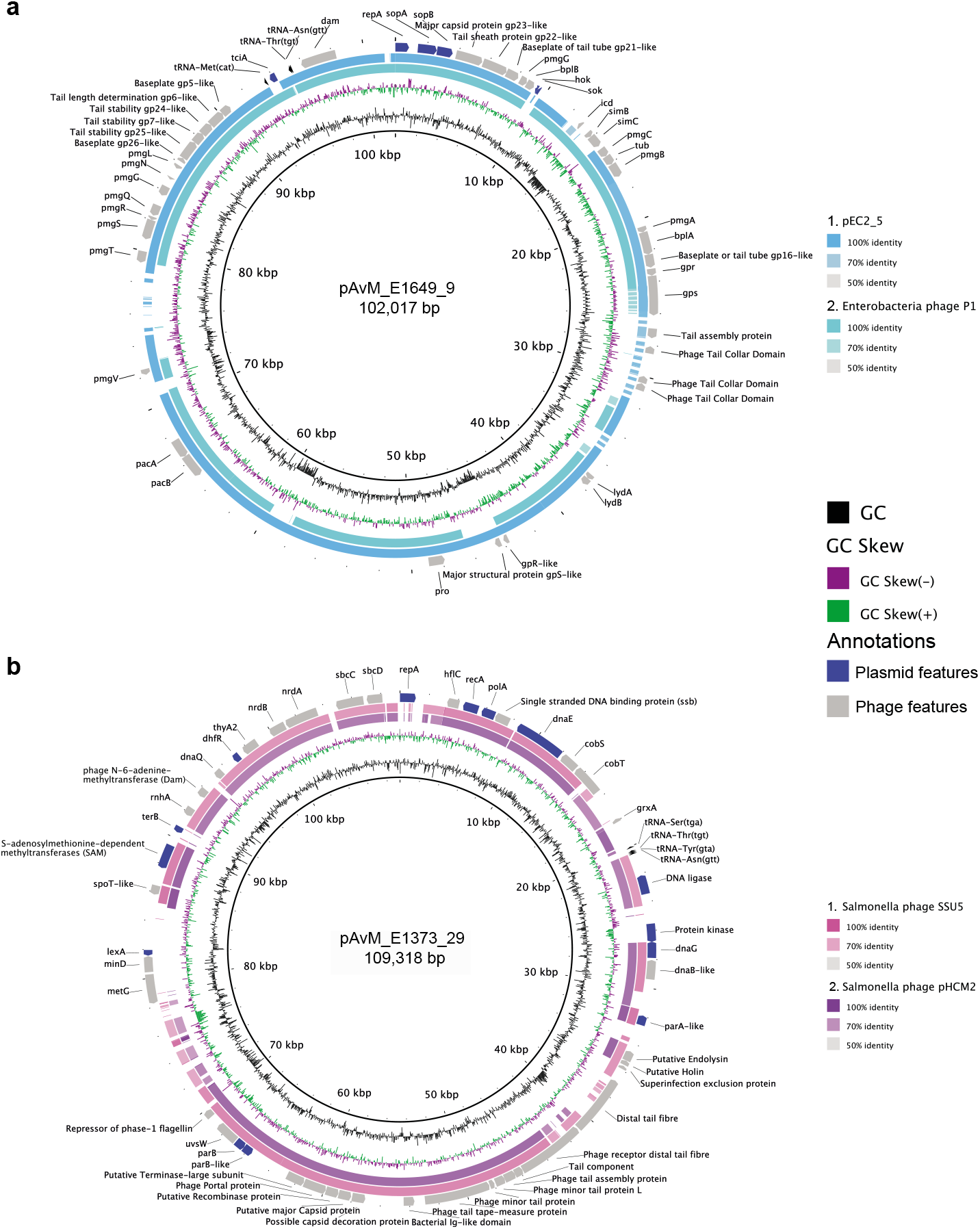
Comparisons between the identified ETEC phage-plasmids and other similar phage-plasmids using blastn. **a)** pAvM_E1649_9 is a P1-like phage-plasmid here compared to Enterobacteria phage P1 (Escherichia virus P1; NC_005856.1) and pEC2_5 (*E. coli* strain EC2_5; CP041960.1). **b)** pAvM_E2980_29, a phage-plasmid similar to the pHCM2 (Salmonella typhi strain CT18; AL513384.1) and SSU5 (Salmonella phage; JQ965645.1) in Salmonella typhi. Blastn comparisons were made using BRIG [84] (v0.95) with the thresholds indicated to the right of each plasmid comparison. Selected phage and plasmid annotations are shown in the outer ring.

Blastn searches confirmed high similarity (at least 80% at the DNA across much of the sequence) of pAvM_E1373_29 to several phage-like plasmids found in *E. coli* including ETEC O169:H41 isolate F8111-1SC3 [20,55], several *bla*_CTX-M-15_ positive phage-like plasmids (pANCO1, pANCO2 [52] and PV234a), as well as a plasmid found in *E. coli* ST648 from wastewater and ST131 isolate SC367ECC [56]. The P1 phage-like plasmid pAvM_1649_9 is most similar to p1107-99K, pEC2_5 isolated from human urine and p2448-3 from a UPEC ST131 isolate isolated from blood. The similarity is most pronounced at the amino acid level. Conservation and synteny are evident when pAvM_1649_9 is compared to P1 phage.

### Prophages present and their cargo genes

Prophages may insert into chromosomes and bring along genes required for lysogeny and lytic cycles and cargo genes that are often picked up when DNA is compacted into the capsid. Cargo genes can significantly benefit the host bacterium by providing additional elements to defence against phage or immune evasion and finally, environmental survival. PHASTER analyses identified prophages in the chromosomes of all ETEC reference isolates and some of the plasmids (Table S4). Putative tellurite resistance operons in isolates E925, E36, E2980 and E1373 were all located in prophages. In addition, *eatA* (in E925, and E1649) and *estA* (STh) genes (E36) were prophage cargo genes.

Many prophage cargo genes identified in this study have properties related to inhibition of cell division. Among these are a variety of *kil* genes which can enhance host bacterial survival in the presence of some antibiotics [57]. Some genes that are core entities within many prophages, such as *zapA* (from E1779_Pph_6), *dicB* and *dicC* (found in phage E1779_Pph_7), also have similar effects as they can inhibit cell division in the presence of antibiotics which raise the broader question in terms of how they are beneficial to the host bacterium.

A different gene of interest is the *yfdR* gene identified in E1779_Pph_7 (gene E1779_04412). YfdR curtails cellular division by inhibiting DNA replication under stress conditions encountered by the bacterial cell. Similarly, the *iraM* gene located in phage E1441_Pph_2 plays a role in RpoS stability.

OmpX homologs were found in numerous phages in this study. They are trans-membrane located and play a role in virulence as well as antibiotic resistance [58]. PerC is often associated with EPEC plasmids, where it seems to have a regulatory role for the attaching and effacing gene, *eaeA* [59]. Its presence as cargo within an ETEC strain phage, E1779_Pph_7 located on the chromosome, is intriguing. Its ability to regulate other virulence genes is yet to be determined. Within the same phage, a *gntR*-like regulatory gene was identified. This gene plays a role in gluconate utilisation and induction of the Entner-Doudoroff pathway [60].

## Discussion

ETEC strains have previously been shown to fall into globally spread genetically conserved lineages which encompass strains with specific virulence factor profiles [18]. The currently widely used ETEC reference strains H10407 (CFA/I) and E24377A (CS1+CS3) are highly divergent from other strains with the same virulence profile sequenced more recently [18] and highlights the need for relevant and representative ETEC reference strains and genomes. The long-read sequenced isolates presented here comprise complete reference genomes with separate chromosomal and plasmid sequences that allow more detailed studies of ETEC and *E. coli* phylogeny. The reference strains are representative isolates of their respective lineage and cluster phylogenetically together with different ETEC isolates sequenced by several other groups (Figure S10).

Previous studies confirmed that ETEC belongs to lineages that have spread globally. These analyses were mainly dependent on the shared core genome of chromosomal genes while conservation of plasmids was indicated by the association between the plasmid-borne toxin and CFs and lineage [18]. Analysis of the plasmids sequenced in the present study showed that the conservation within ETEC lineages also include plasmids.

Blast analyses confirm that the plasmids identified in this study are often highly homologous to other plasmids identified by either long-read or Illumina sequencing present in GenBank. For instance, the 94.5 kb plasmid pAvM_1441_18 was 98% identical to two 96 kb and 82 kb plasmids belonging to ETEC O25: H16 isolates ATCC 43886/E2539C1 and 2014EL-1346-6 sequenced by PacBio by Smith *et al*., [20] (Figure 1e and Figure S6). Plasmid pAvM_E1441_18 is the major virulence plasmid of this lineage carrying genes encoding LT and CS6.

The larger plasmid in E1441 (pAvM_E1441_17) carries both the genes for ETEC CF CS21 and antibiotic resistance determinants. Furthermore, complete conjugation machinery was present suggesting that this is most likely a self-transmissible plasmid, though this was not confirmed. Movement of such a plasmid would result in the spread of ETEC virulence genes and AMR determinants.

Interestingly, Wachsmuth et al., [44] analysed transfer frequencies in ETEC O25:H16 isolates (the same serogroup was identified in E1441) and found evidence that resistance to tetracycline and sulfathiazole was transferred but not the genes encoding LT [44]. The same study found evidence of two large plasmids of similar size [44] corroborating our findings of two plasmids of similar size in E1441, one with *eltAB* and *cssABCD* without the *tra*-operon (pAvM_E1441_18) and the other putatively mobile plasmid (pAvM_E1441_17) carrying the *sul1* and *tet*(A) genes as well as the *lng* operon encoding CF CS21. Since ATCC 43886/E2539C1, E1441 and 2014EL-1346-6, have been isolated in the 1970s, 1997, and 2014, respectively, our findings indicate that E1441 represent an ETEC lineage with stable plasmid content and putative ability to transfer antibiotic resistance and the CS21 operon by transfer of one of the plasmids. Furthermore, pAvM_E1441_17 is a multi-replicon plasmid. Multi-replicon plasmids have been described as a way to broaden their host range, i.e. possibility to be transferred between bacteria of different phylogenetic groups [61,62]. Whether this plasmid type is found in other *E. coli* remains to be investigated but the finding that the L4 lineage retains both plasmids in isolates collected over time and worldwide indicate a strong selective force to keep the extra-chromosomal contents of both plasmids.

The ETEC O169:H41 isolate F8111-1SC3 plasmid unnamed 2 [20,55] is highly similar to pAvM_E1373_28 (Figure 1h and Figure S9). The F8111-ISC3 isolate is part of a CDC collection of ETEC isolates from cruise ship outbreaks and diarrheal cases in US 1996-2003. The antibiotic resistance profiles of these isolates were determined [55] and most isolates of O group 169 were tetracycline resistant consistent with the findings of the *tet* gene in E1373 isolated in Indonesia in 1996. ETEC diarrhoea caused by O169:H41 and STp CS6 isolates is repeatedly reported to cause diarrhoea, particularly in Latin America [45,63–65]. Among the cruise ship isolates is the sequenced and characterised virulence plasmid pEntYN10 encoding STp and CS6, described as unstable and easily lost in vitro [63,66]. The E1373 plasmid; AvM_E1373_28 is highly homologous to pEntYN10 (Figure 1h and Figure S9) and the virulence profile of ETEC O169: H41 is conserved in isolates collected globally. Hence, the instability of the plasmid is incongruent with current data indicating that plasmids are stable within this lineage and serotype.

Interestingly, two distinctive extra-chromosomal elements which are highly similar to P1 and SSU5 phage were identified among the 8 ETEC reference strains sequenced (Figure 2, Figure S14 and Table S4). The SSU5-like element carries several genes that allow it to be functional as a plasmid and belongs to the pHCM2-like family of Phage-Plasmids (Figure 2b) [49]. These plasmids are devoid of virulence factors, transposons and antibiotic markers but, they contain a significant number of DNA metabolism and biosynthesis genes and they may contain bacteriophage inhibitory genes that have not yet been identified. Interestingly, several SSU5 phage-like plasmids have been shown to carry the ESBL gene *bla*_CTX-M15_ in extra-intestinal pathogenic *E. coli* isolates [51]. ESBL resistance seems to be absent or low in ETEC and the SSU5 phage-like plasmid pAvM_E1373_29 does not contain antibiotic resistance genes. A recent study investigating the distribution of phage-plasmids show that the phage homologs tend to be more conserved and the plasmid homologs more variable [67]. This is also seen in the phage-plasmids identified here, e.g., genes that could be advantageous to the host cell linked to metabolism and biosynthesis.

### Conclusion

We provide fully assembled chromosomes and plasmids with manually curated annotations that will serve as new ETEC reference genomes. The in-depth analysis of gene content, synteny and correct annotations of plasmids will also help to elucidate other plasmids with and without virulence factors in related bacterial species. The ETEC reference genomes compared to other long-read sequenced ETEC genomes confirm that the major ETEC lineages harbour conserved plasmids that have been associated with their respective background genomes for decades. This confirms that the plasmids and chromosomes of ETEC are both crucial for ETEC virulence and success as pathogens.

## Methods

### Selection of strains

Initially one to two ETEC strains within each of the lineage (L1-L7)-specific CF profile were chosen from the University of Gothenburg large collection of ETEC strains [18] for PacBio sequencing. The strains were selected based on the location and year of isolation to represent strains isolated from patients with diarrhoea from diverse geographical locations and at different time-points. After the genomes had been sequenced, assembled, circularised and annotated a second selection was made for manual curation of the genomes. This selection was made based on the quality of the genome assembly and the circularisation. The whole genomes of the ETEC reference strains were compared with one or two other long-read sequenced ETEC strains belonging to the same lineage by progressiveMAUVE and showed that the strains are colinear (Figure S15). One representative ETEC genome from each lineage was annotated, with emphasis on the plasmids. The physical ETEC reference strains are available upon request.

### Phenotypic toxin and CF analyses

ETEC isolates were identified by culture on MacConkey agar followed by an analysis of LT and ST toxin expression using GM1 ELISAs [43]. The expression of the different CFs was confirmed by dot-blot analysis [43]. Isolates had been kept in glycerol stocks at −70 °C, and each strain has been passaged as few times as possible.

### Antibiotic susceptibility testing

All ETEC isolates were tested against 14 antimicrobial agents and their minimum inhibitory concentration was determined by broth microdilution using EUCAST methodology [47]. The antimicrobial agents were: ampicillin, amoxicillin-clavulanic, oxacillin, ceftazidime, ceftriaxone, doxycycline, tetracycline, nalidixic acid, norfloxacin, azithromycin, erythromycin, chloramphenicol, nitrofurantoin and sulfamethoxazole-trimethoprim. All antibiotics were purchased from Sigma-Aldrich. The *E. coli* ATCC 25922 was used as quality control. The MIC was recorded visually as the lowest concentration of antibiotic that completely inhibits growth.

### DNA extraction and sequencing

Strains from each lineage (L1-L7) were SMRT-sequenced on the PacBio RSII. A hybrid *de novo* assembly was performed combining the reads from both the SMRT-sequenced and Illumina sequenced strains.

For Single-Molecule Real-Time (SMRT) sequencing (Pacific Bioscience) long intact strands of DNA are required. The genomic DNA extraction was performed as follows.

Isolates were cultured in CFA broth overnight at 37°C followed by cell suspension in TE buffer (10 mM Tris and 1 mM EDTA pH 8.0) with 25% sucrose (Sigma) followed by lysis using 10 mg/ml lysozyme (in 0.25 Tris pH 8.0) (Roche). Cell membranes were digested with Proteinase K (Roche) and Sarkosyl NL-30 (Sigma) in the presence of EDTA. RNase A (Roche) was added to remove RNA molecules. A phenol-chloroform extraction was performed using a mixture of Phenol:Chloroform:Isoamyl Alcohol (25:24:1) (Sigma) in phase lock tubes (5prime). To precipitate the DNA 2.5 volumes 99% ethanol and 0.1 volume 3 M NaAc pH 5.2 was used followed by re-hydration in 10 mM Tris pH 8.0. DNA concentration was measured using NanoDrop spectrophotometer (NanoDrop). On average 10 μg for PacBio sequencing. Library preparation for SMRT sequencing was prepared according to the manufacturers’ (Pacific Biosciences) protocol. The DNA was stored in E buffer and sequenced at the Wellcome Sanger Institute. Isolates were sequenced with a single SMRTcell using the P6-C4 chemistry, to a target coverage of 40–60X using the PacBio RSII sequencer.

### Assembly

The resulting raw sequencing data from SMRT sequencing were *de novo* assembled using the PacBio SMRT analysis pipeline (https://github.com/PacificBiosciences/SMRT-Analysis) (v2.3.0) utilising the Hierarchical Genome Assembly Process (HGAP) [68]. For all samples, the unfinished assembly produced a single, non-circular, chromosome plus some small contigs, some of which were plasmids or unresolved assembly variants. Using Circlator [69] (v1.1.0), small self-contained contigs in the unfinished assembly were identified and removed, with the remaining contigs circularised. Quiver [68] was then used to correct errors in the circularised region by mapping corrected reads back to the circularised assembly. As the strains had also been short read sequenced, and this data is of higher base quality, the short reads from the Illumina sequencing were used in combination with the long reads using Unicycler [70] to generate high-quality assemblies.

Fully circularised chromosomes and plasmids were achieved for the majority of the strains. Cross-validation of the assemblies was performed where two or three strains of a lineage were sequenced (Figure S15). A single assembly from each lineage was chosen to act as the representative reference genome, with priority given to assemblies with the most complete and circularised chromosome and plasmids. In total, one chromosome and 5 out of the 29 plasmids could not be circularised (independent on the two strains that were sequenced initially) out of the 8 selected representative strains. These are indicated in Table 2 and Table S1. Between two and five plasmids were identified in the eight strains. Shorter contigs that could not be assembled properly contained phage genes and are included in the genomes and annotated as prophages Table S4). Socru was used to validate the assembly of the chromosome, they all have biologically valid orientation and order of rRNA operons with a type GS1.0, which is seen in most *E. coli* in the public domain [71].

### Phylogenetic tree

The phylogenetic relationship between the ETEC reference genomes to other ETEC and *E. coli* commensals and pathotypes was investigated. The following collections were included: ETEC-362 [18], ECOR [72] and the Horesh collection [73] along with additional ETEC genomes from several studies [20,24,26,27,45,74,75]. The reads of identified ETEC genomes from other studies were downloaded from GenBank and assembled using Velvet. Long-read sequenced ETEC genomes were included in the tree and were not re-assembled. The phylogroup of the ETEC strains was determined using ClermonTyping [32] (v20.03). The virulence profile of the ETEC strains was determined using ARIBA [76] (v2.14.16) with default settings using the custom ETEC virulence database (https://github.com/avonm/ETEC_vir_db). A total of 1,066 genomes was included in the phylogenetic tree. The alignment of core genes (n = 2,895) identified by Roary [77] (v3.12.0) was converted to a SNP-only alignment using snp-sites [78]. A phylogenetic tree was produced with IQ-TREE [79] (v1.6.10) using a GTR gamma model (GTR+F+I) optimised using the built-in model test and visualised using the R package ggtree [80].

### Gene prediction, annotation and comparative analysis

The final assembly was annotated using Prokka [81] (v1.14.6). The annotations of all plasmids generated by Prokka were manually checked using the genome viewer Artemis [82] and Geneious 2018.11.1.5 (http://www.geneious.com) together with blastp. Annotations of known ETEC virulence genes (colonisation factors, toxins, *eatA* and *etpBAC*) were added after blast+ [83] analysis using the reference genes available in the ETEC virulence database (https://github.com/avonm/ETEC_vir_db) and their annotations updated accordingly. The LT and ST alleles were determined according to Joffre et al., (https://github.com/avonm/ETEC_toxin_variants_db) [15,17]. Where required, PFAM domains were searched using jackhammer to back up any identified protein using blastp (https://www.ebi.ac.uk/Tools/hmmer/search/jackhmmer). Blastn and tblastx were used for plasmid comparison, using both NCBI website or within BLAST Ring Image Generator (BRIG) [84] (v0.95).

### Incompatibility groups

Due to the discrepancy in databases two approaches was used to determine the Inc groups of the 25 plasmids. PlasmidFinder was used with a threshold for minimum % identity at 95% and minimum coverage of 60%. The plasmids were further characterised by pMLST [34], except for IncY which are a group of prophages that replicate in a similar manner as autonomous plasmids (Additional File 3). IncB/O/K/Z plasmids were further typed by blastn comparison to the reference B/O (M93062), K (M93063) and Z (M93064) replicons.

### oriT prediction

The location of the *oriT* in the plasmids, if present, was predicted using oriTFinder [85] with Blast E-value cut-off set to 0.01.

### Genomic antibiotic resistance profiling

The identification of antibiotic resistance genes, located on both the chromosome and plasmid(s) as well as the presence of efflux pumps and porins known to confer resistance to antibiotics. The results were obtained by running ARIBA [76] using the CARD database [86] with the default settings (minimum 90% sequence identity and no length cut-off). ARIBA combines a mapping/alignment and targeted local assembly approach to identify AMR genes and variants efficiently and accurately from paired sequencing reads. The heatmaps were visualised using Phandango [87]. The presence of chromosomal mutations in *gyrA* and *parC* was determined with ResFinder (v3.2) from the Center of Genomic Epidemiology [88].

### Virulence gene prediction

The ETEC assemblies from the ETEC-NCBI collection (Additional file 4) were screened using abricate [89] with default settings against the ETEC virulence database (https://github.com/avonm/ETEC_vir_db) for virulence gene (including *eatA* and *etpBAC*) predication. A subset of the isolates in the ETEC-NCBI dataset have previously been analysed for the presence of EatA where a sample with negative PCR but positive western blots were included as positive [74]. Here, only isolates harbouring the *eatA* and *etpBAC* genes are considered positive.

### Prophage prediction

The complete FASTA sequence of each ETEC reference genome was searched for phage genes and prophages using PHASTER (phaster.ca) [90]. The identified intact prophages are listed in Table S4. All prophage contained cargo genes but only recognisable genes are stated, not any hypothetical. Additional questionable and not intact prophages were identified but have not been included here. The prophages have been given a specific identifier name and are also annotated as a mobile_element in the submitted chromosome and or plasmid(s) of each strain.

### Insertion sequences

Insertion sequences in the plasmids as well as surrounding the CS2 loci located on the chromosome of E1649 were annotated using both Galileo AMR software [91] and the ISFinder database [92]. Complete and partial IS elements were annotated (>95% identity with hits in ISFinder) along with the present genes encoding transposases. Three new insertion sequences were detected in this analysis and were submitted to ISFinder as TnEc2, TnEc3 and TnEc4. Transposons and other mobile elements (integrons and group II introns) were also identified using Galileo AMR and blastn against public databases.

## Supplementary information

**Additional file 1:** Supplemental Figures and Tables

**Additional file 2:** Detailed description of ETEC reference plasmids

**Additional file 3:** Excel file with plasmid classification – Inc groups

**Additional file 4:** Excel file with metadata of ETEC and *E. coli* genomes included in the phylogenetic tree

**Additional file 5:** Excel file with a compiled list of all accession numbers for chromosomes and plasmids

## Acknowledgements

No acknowledgements to mention.

## Authors’ contributions

AvM conceived and designed the experiments, performed the experiments, analysed the data, contributed reagents/materials/analysis tools, prepared figures and/or tables, authored the paper and approved the final draft.

GB and AvM annotated all IS elements, transposons as well as other mobile elements, contributed to the paper and approved the final draft.

DP and AvM annotated the identified prophages, contributed to the paper and approved the final draft.

CB performed the *in silico* analysis of the genomic antibiotic resistance profiling, contributed to the paper and approved the final draft.

EJ performed the antibiotic resistance profiling, contributed to the paper and approved the final draft.

AJP assembled the genomes, contributed to the paper and approved the final draft.

AMS conceived and designed the experiments contributed to the paper and approved the final draft.

GD conceived and designed the experiments and approved the final draft.

ÅS conceived and designed the experiments, analysed data, authored the paper and approved the final draft.

## Funding

AvM, AMS and ÅS were supported by the Swedish Foundation for Strategic Research (grant nr. SB12-0072). AvM was also supported by The Swedish Research Council (grant nr. 2018-06828) and the Swedish Society for Medical Research (P18-0140). AJP was supported by the Biotechnology and Biological Sciences Research Council (BBSRC); this research was funded by the BBSRC Institute Strategic Programme Microbes in the Food Chain BB/R012504/1. GD was supported by the Wellcome Trust (grant WT 098051).

## Availability of data

The datasets supporting the conclusions of this article are included within the articles and its additional files. The sequencing data generated in this study has been submitted to EMBL (Additional file 4 and 5). The physical ETEC reference strains can be requested by contacting the corresponding author Astrid von Mentzer. The database used for annotating ETEC virulence factors, ETEC virulence database, including the LT and ST alleles can be found in the github repositories: https://github.com/avonm/ETEC_vir_db and https://github.com/avonm/ETEC_toxin_variants_db.

An interactive version of the core genome phylogeny of the 1,065 *E. coli* and ETEC isolates along with the ETEC reference strains (Figure S10) reported here is accessible at https://microreact.org/project/2ZZzaHzeXbMEw9U2MAk7pK?tt=cr

## Availability of material

Requests for obtaining clinical isolates collected as part of this study should be addressed to the corresponding author. Exchange of clinical isolates should always be in agreement with the University of Gothenburg.

## Ethics approval and consent to participate

Not applicable.

## Consent for publication

Not applicable.

## Competing interests

The authors declare that there are no conflicts of interest.

## Notes

https://github.com/avonm/ETEC_toxin_variants_db

https://github.com/avonm/ETEC_vir_db

https://microreact.org/project/2ZZzaHzeXbMEw9U2MAk7pK?tt=cr

### Competing Interest Statement

The authors have declared no competing interest.

### Summary of Updates

Github repositories have been added for easy access to databases used for virulence gene analysis. A phylogenetic tree showing ETEC references in context with additional ETEC genomes and E. coli from two collections (Figure S10 and S11). An interactive tree is also available in microreact and the metadata can be downloaded within microreact. Additional metadata of the isolates included in the phylogenetic tree is available in an excel file (Additional File 4) Additional plasmid maps showing additional annotations are included in Additional File 1 (Figure S2-S9).

