## Additional file 1: Supplementary figures and tables for "Long-read-sequenced reference genomes of the seven major lineages of enterotoxigenic *Escherichia coli* (ETEC) circulating in modern time"

### Additional file 1
### Supplementary Figures and Tables

Figures S1-S16

Tables S1-S4

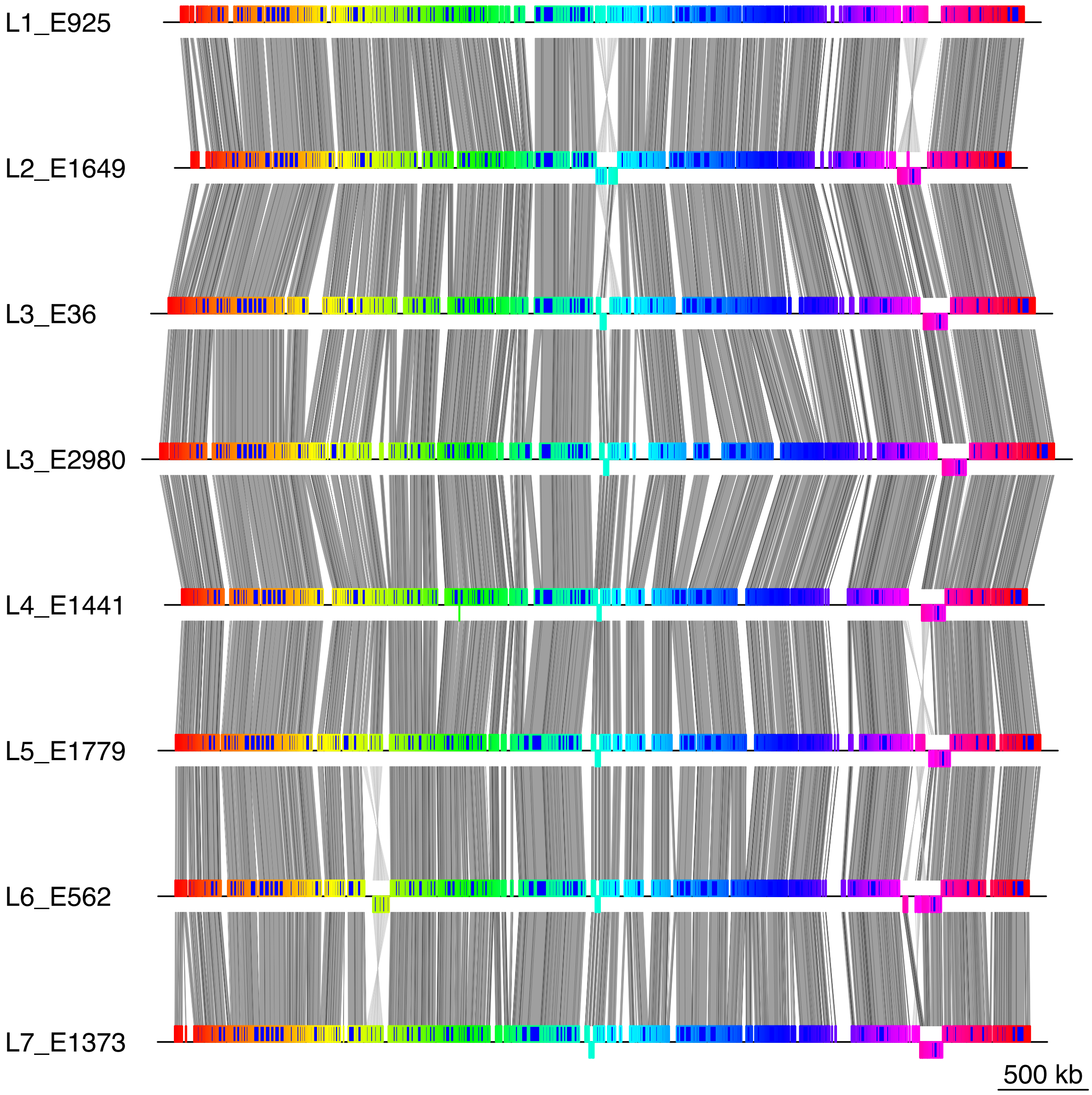


**Figure S1:** Comparison of the chromosomes of each ETEC reference genome using progressiveMAUVE. Circularized chromosomes extracted from the reference ETEC strains of lineage 1 through 7 (two reference strains belong to L3, one is CFA/I and the other CS7). The chromosomes are to a large extent colinear. Visualized using genoplotR.

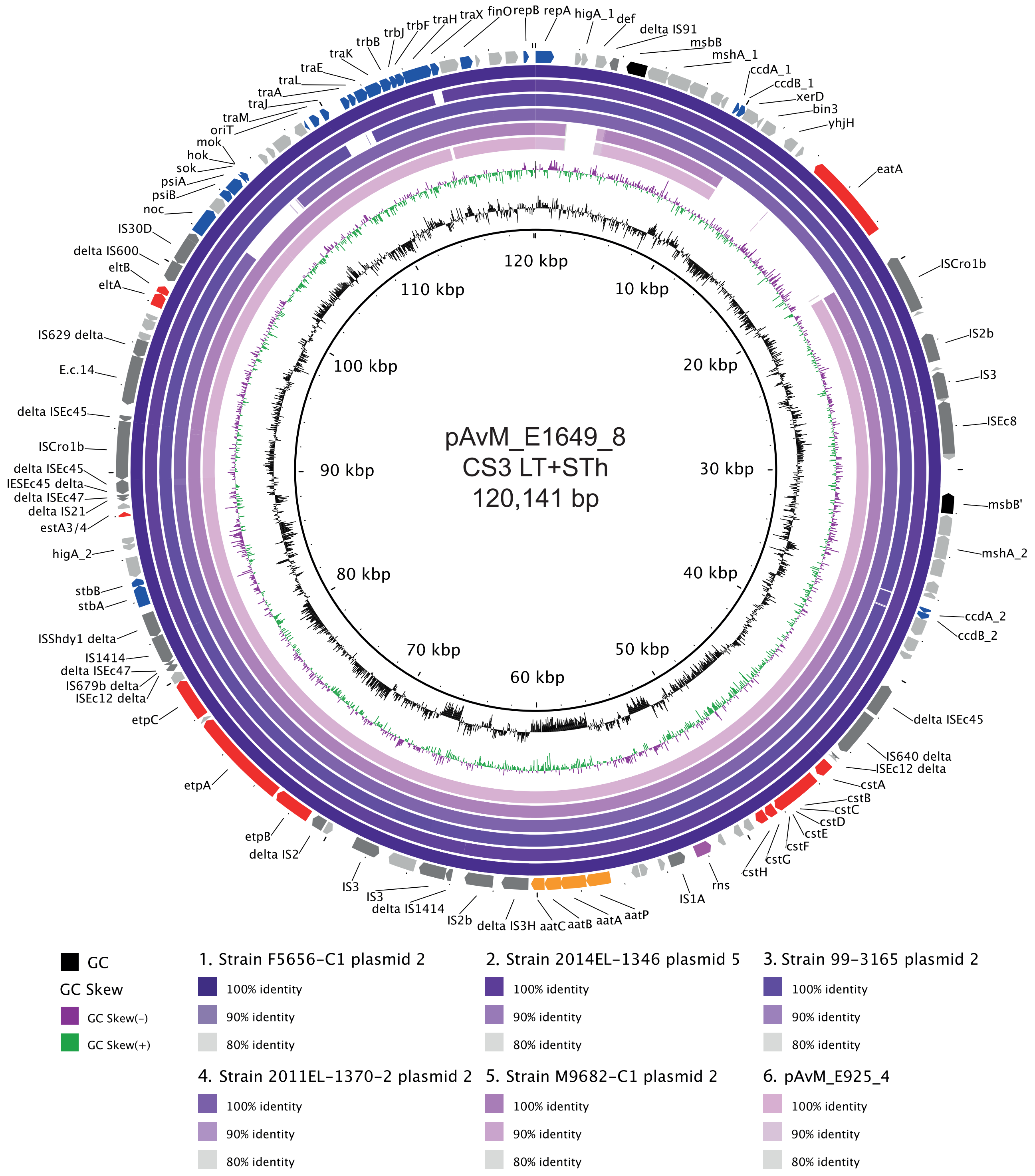


**Figure S2:** Comparison between the ETEC reference plasmids harbouring colonisation factors and other PacBio-sequenced ETEC plasmids using blastn. Plasmid pAvM_E1649_8 (CS3) as reference and pAvM_925_4 (CS3) compared to the following ETEC plasmids: F5656-C1 plasmid 2 (USA, CP024262.1), 2014-EL-1346-6 plasmid 5 (2014, USA; CP024237.1), 99-3165 plasmid 2 (USA; CP029980.1), 2011EL-1370-2 plasmid 2 (2011, USA; CP022914.1) and M9682-C1 plasmid 2 (1975, USA; CP024277.1).

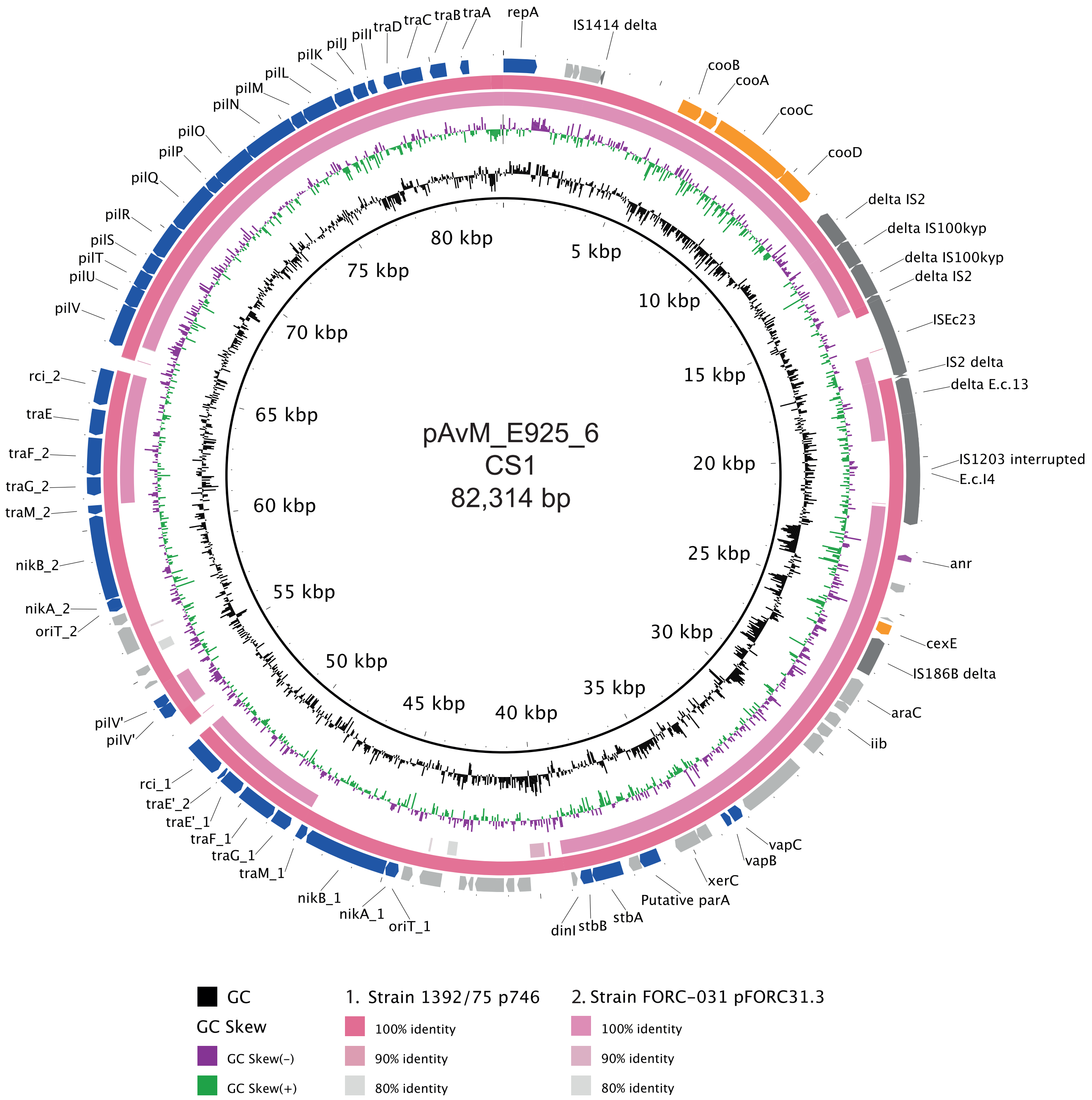


**Figure S3:** Comparison between the ETEC reference plasmids harbouring colonisation factors and other PacBio-sequenced ETEC plasmids using blastn. Plasmid pAvM_E925_6 (CS1) compared to ETEC plasmids 1392/75 p746 (1973; FN822748.1) and pFORC31.3 (2004, Korea; CP013193.1).

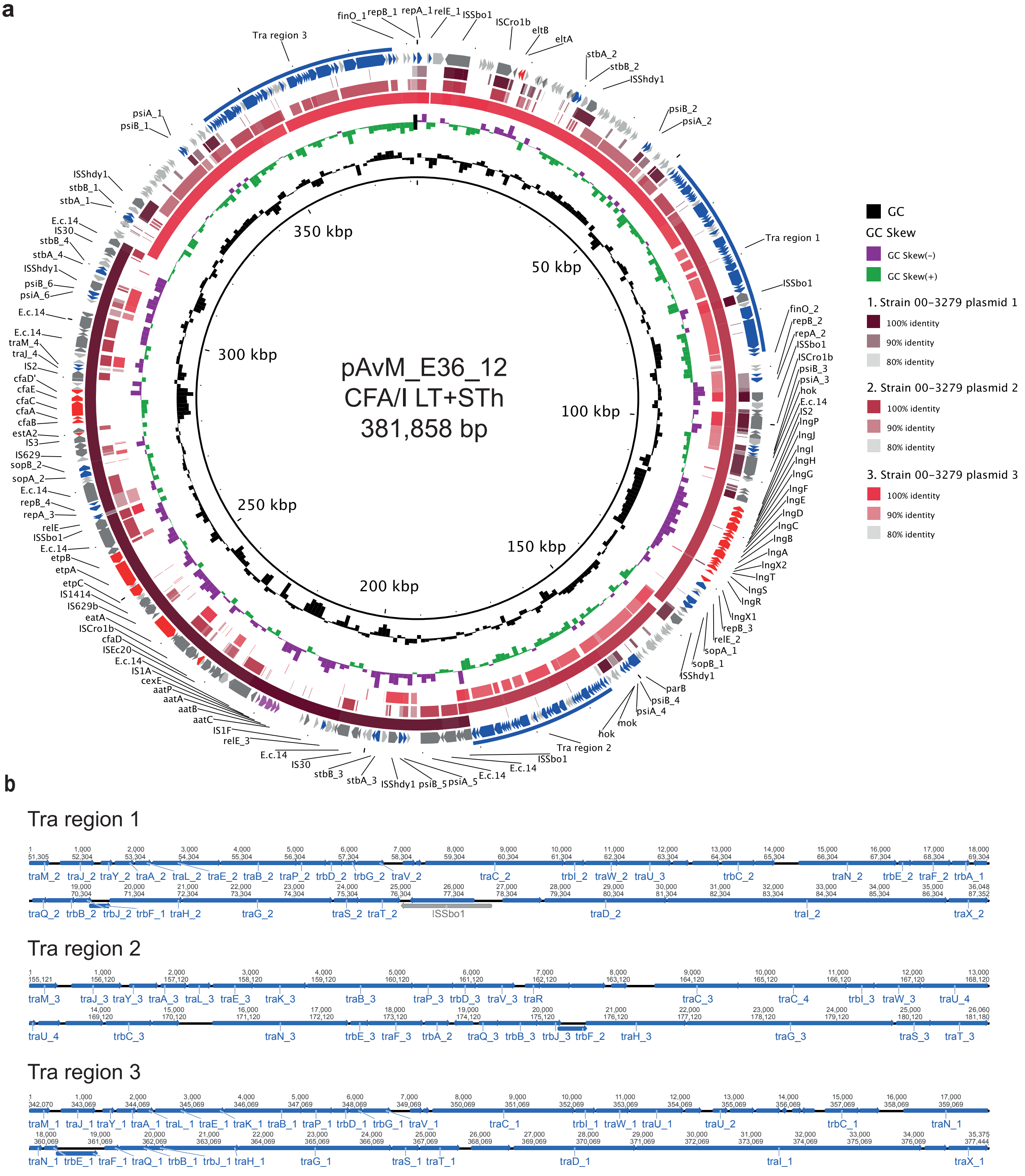


**Figure S4:** Comparison between the ETEC reference plasmids harbouring colonisation factors and other PacBio-sequenced ETEC plasmids using blastn. **a)** pAvM_E36_12 (CFA/I) compared to plasmids 1-3 (p1: CP024294.1; p2 CP024295.1; p3: CP024296.1) from ETEC strain 00-3279 (USA). **b)** Organisation of Tra regions 1, 2 and 3.

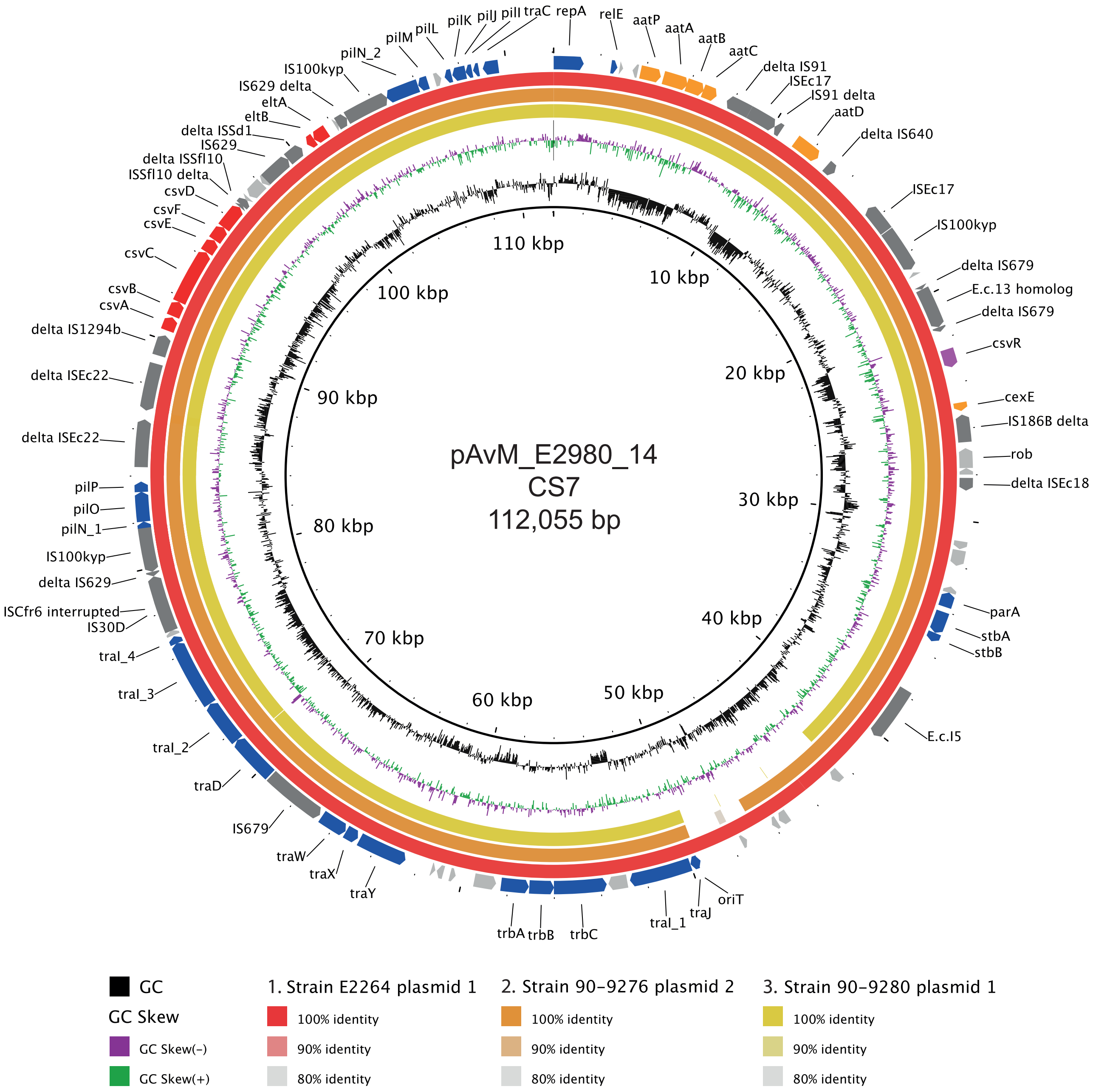


**Figure S5:** Comparison between the ETEC reference plasmids harbouring colonisation factors and other PacBio-sequenced ETEC plasmids using blastn. Plasmid pAvM_E2980_14 (CS7) compared to E2264 plasmid 1 (2006, Bangladesh; CP023350.1), 90-9276 plasmid 2 (1988, Bangladesh; CP024298.1) and 90-9280 plasmid 1 (1988, Bangladesh; CP024241.1).

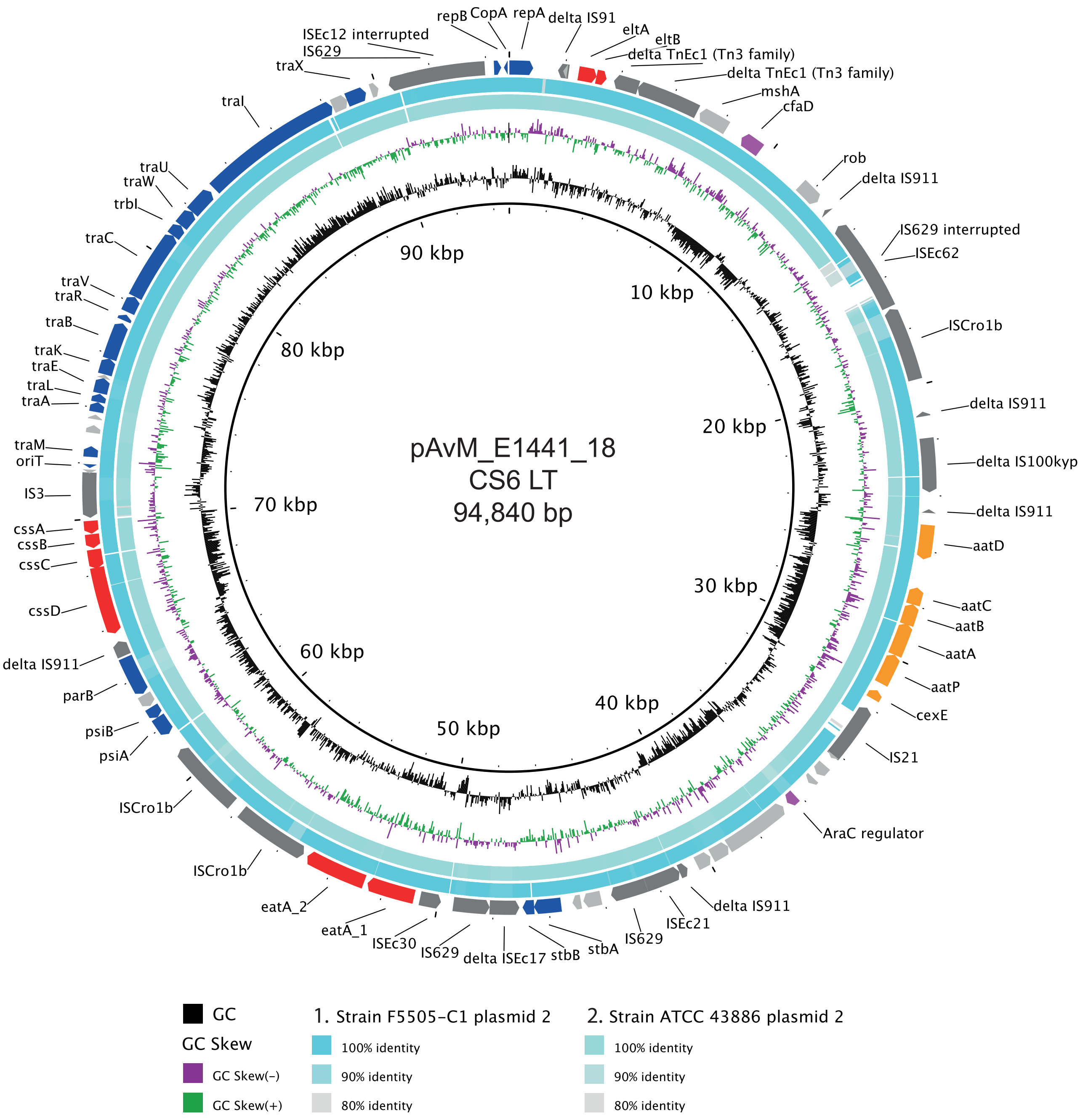


**Figure S6:** Comparison between the ETEC reference plasmids harbouring colonisation factors and other PacBio-sequenced ETEC plasmids using blastn. pAvM_E1441_18 (CS6) compared to F5505-C1 plasmid 2 (2013, Sweden; CP023259.1) and ATCC 43886 plasmid 2 (CP024255.1).

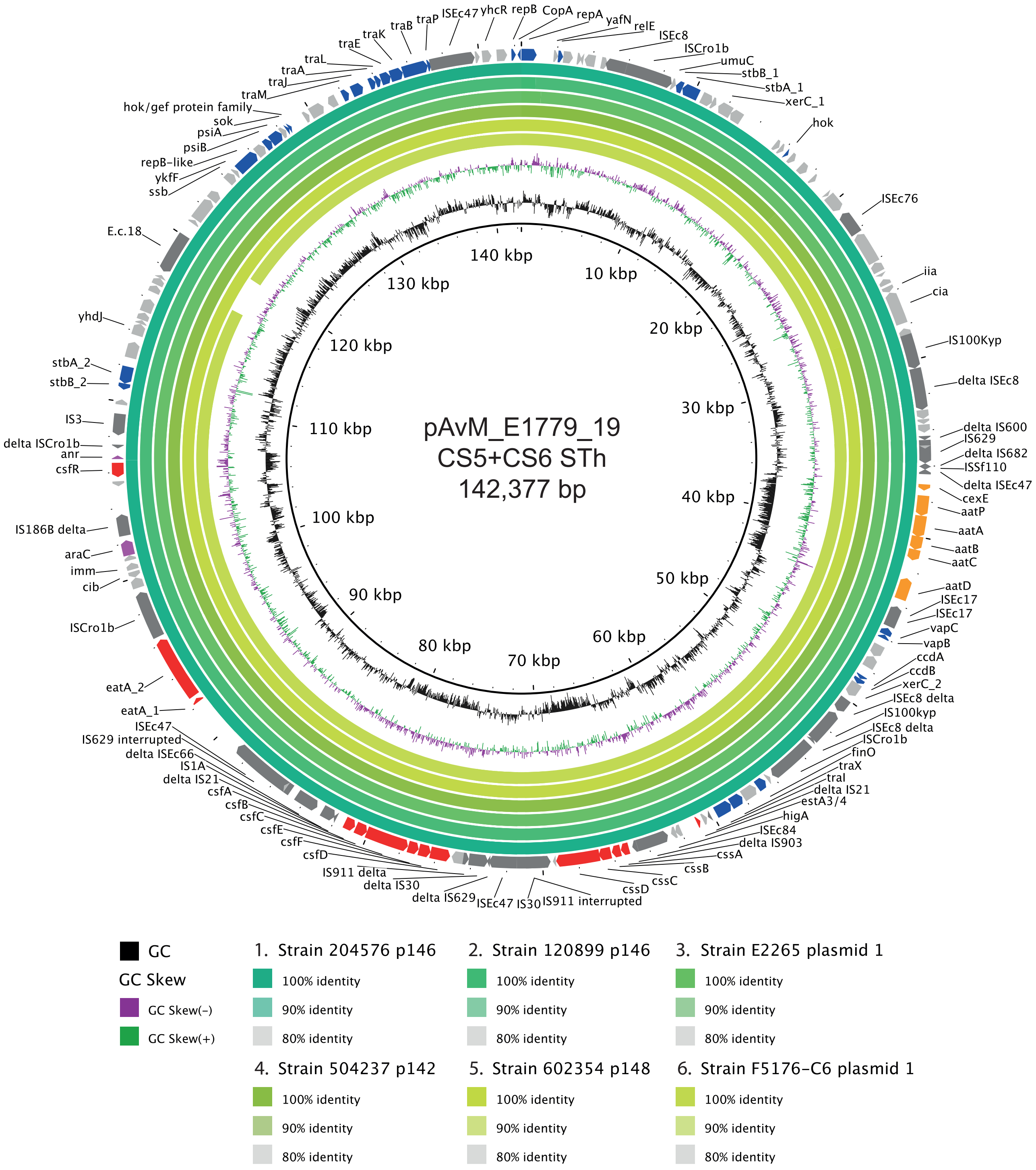


**Figure S7:** Comparison between the ETEC reference plasmids harbouring colonisation factors and other PacBio-sequenced ETEC plasmids using blastn. pAvM_E1779_19 (CS5+CS6) compared to 204576 p146 (2010, Mali; CP025908.1), 120899 p146 (2012, Gambia; CP025917.1), E2265 plasmid 1 (2006, Bangladesh; CP023347.1), 504237 p142 (2010, India; CP025863.1), 602354 p148 (2009, Bangladesh; CP025848.1) and F5176-C6 plasmid 1 (1997; CP024668.1).

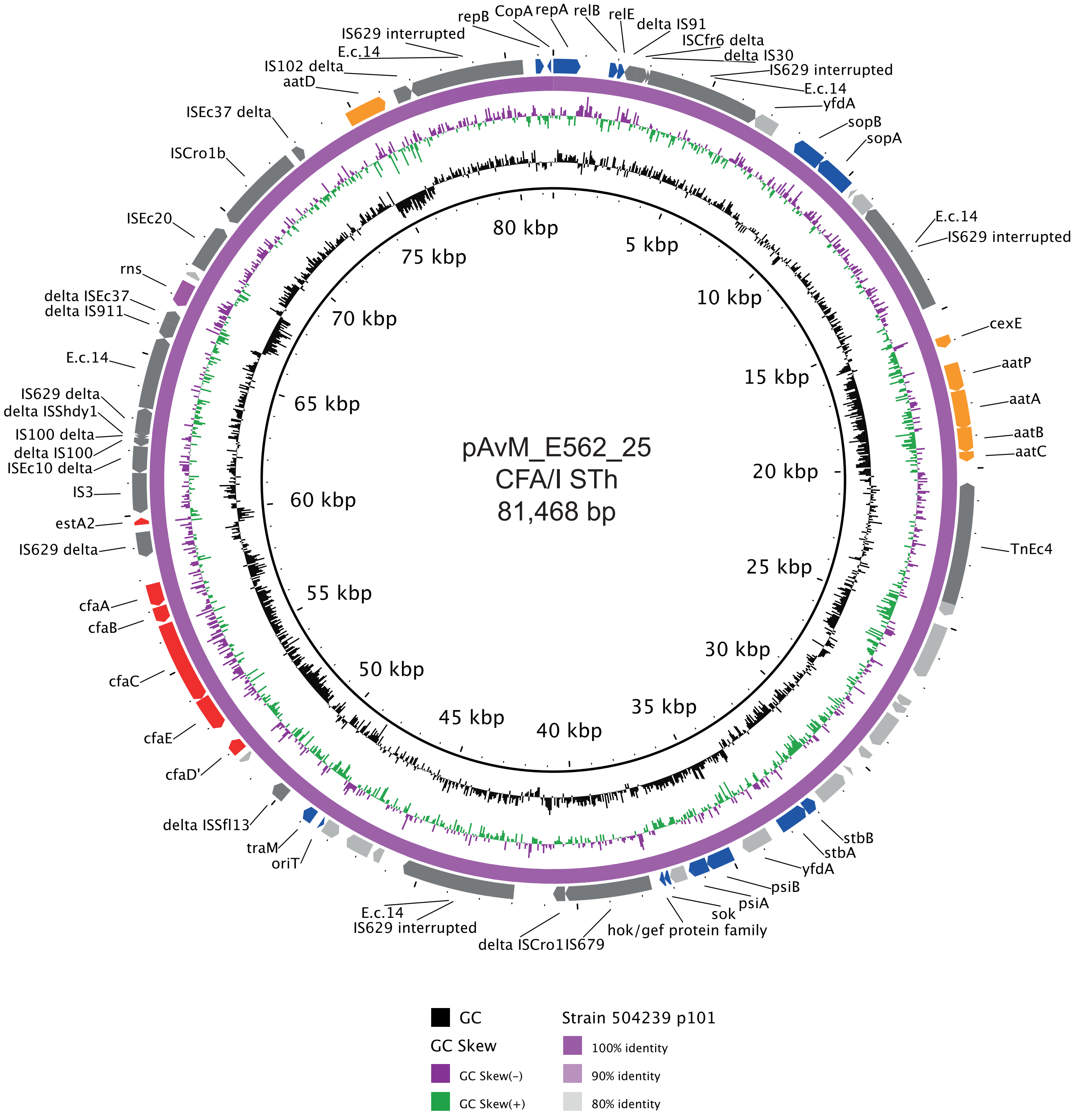


**Figure S8:** Comparison between the ETEC reference plasmids harbouring colonisation factors and other PacBio-sequenced ETEC plasmids using blastn. pAvM_E562_25 (CFA/I) compared to p504239_101 (2010, India; CP025860.1).

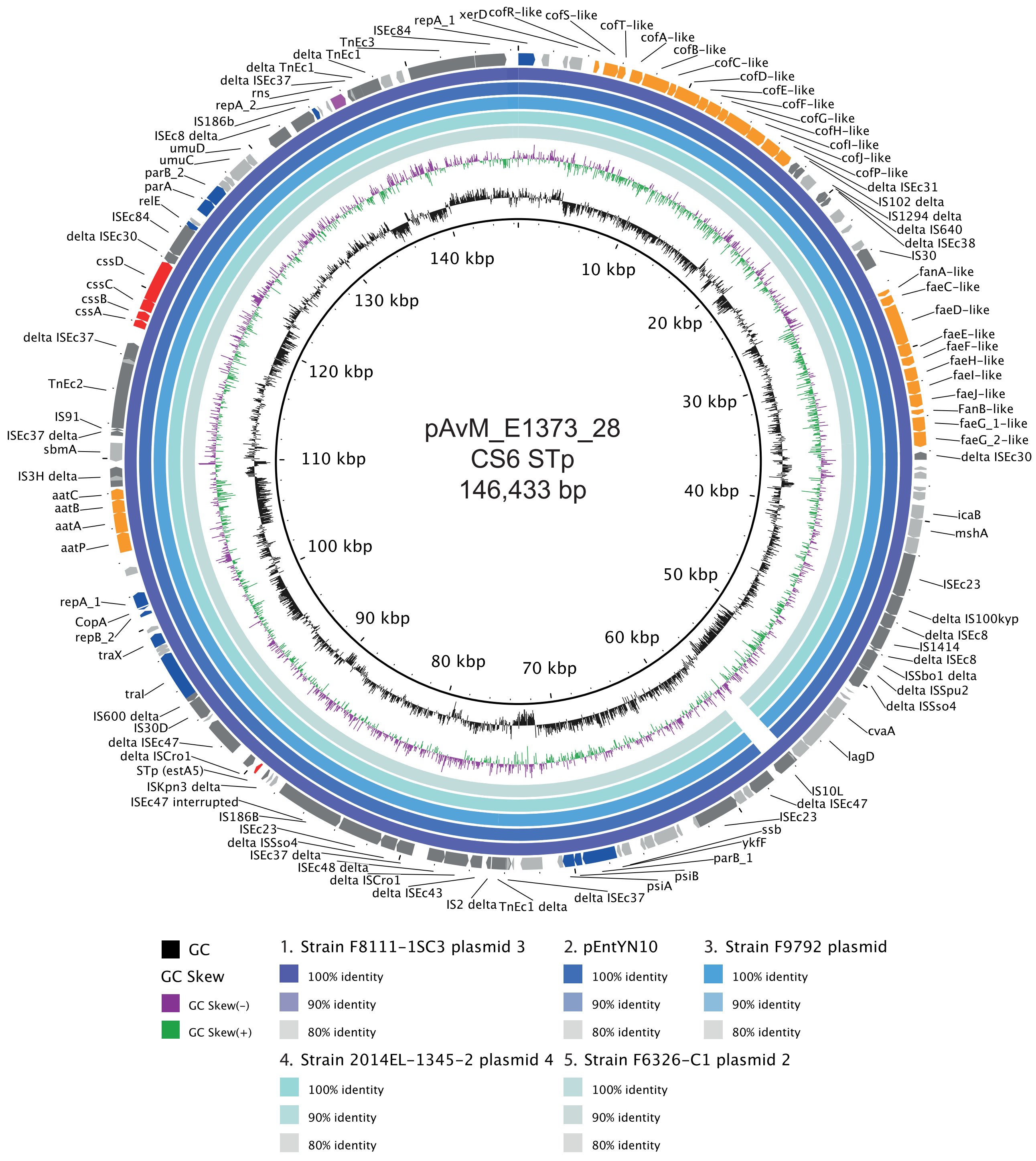

**Figure S9:** Comparison between the ETEC reference plasmids harbouring colonisation factors and other PacBio-sequenced ETEC plasmids using blastn. pAvM_E1373_28 (CS6) compared to F8111-1SC3 plasmid 3 (USA; CP024272.1), pEntYN10 (1991, Japan; AP014654.2), F9792 plasmid (USA; CP023274.1), 2014EL-1345-2 plasmid 4 (2014, USA; CP024227.1) and F6326-C1 plasmid 2 (1998, USA; CP024265.1).

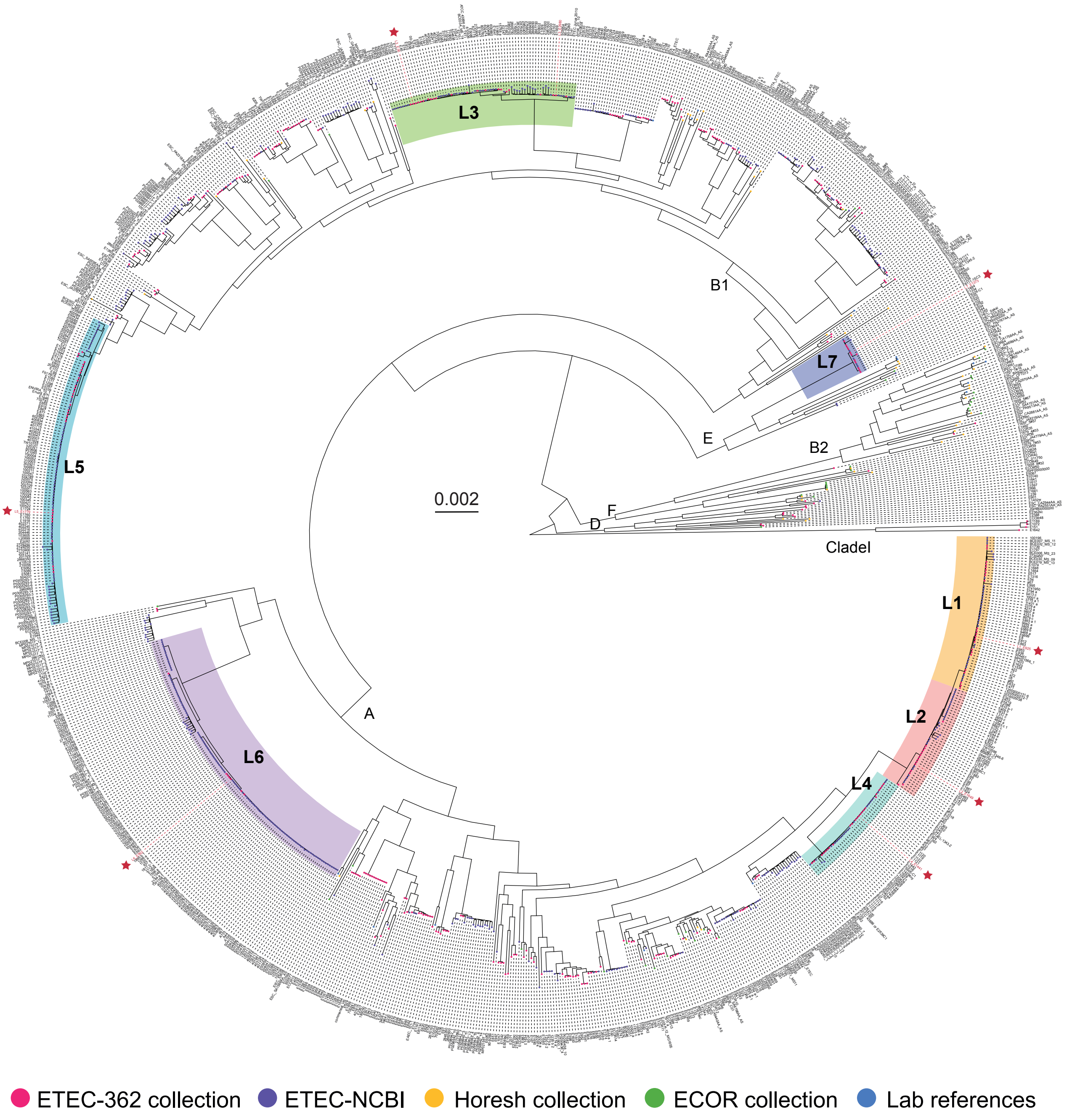


**Figure S10:** Phylogenetic analysis of ETEC genomes in context of *E. coli* genomes. Phylogenetic tree and network depicting the relationship of genomes from multiple ETEC studies (refs. 20, 24, 26, 27, 45, 76, 77) and representative *E. coli* genomes including *E. coli* commensals and pathotypes (refs. 74 and 75). Maximum likelihood phylogeny (1000 ultra fast parametric bootstraps, midpoint rooted). Strain IDs are located on the tips and the ETEC reference strains are colored in red and marked with a red star. The phylogroups are indicated in the tree and the major lineages presented in ref. 18 are highlighted. The scale bar indicates the number of substitutions per site. A tree with bootstrap values is provided in Figure S11.

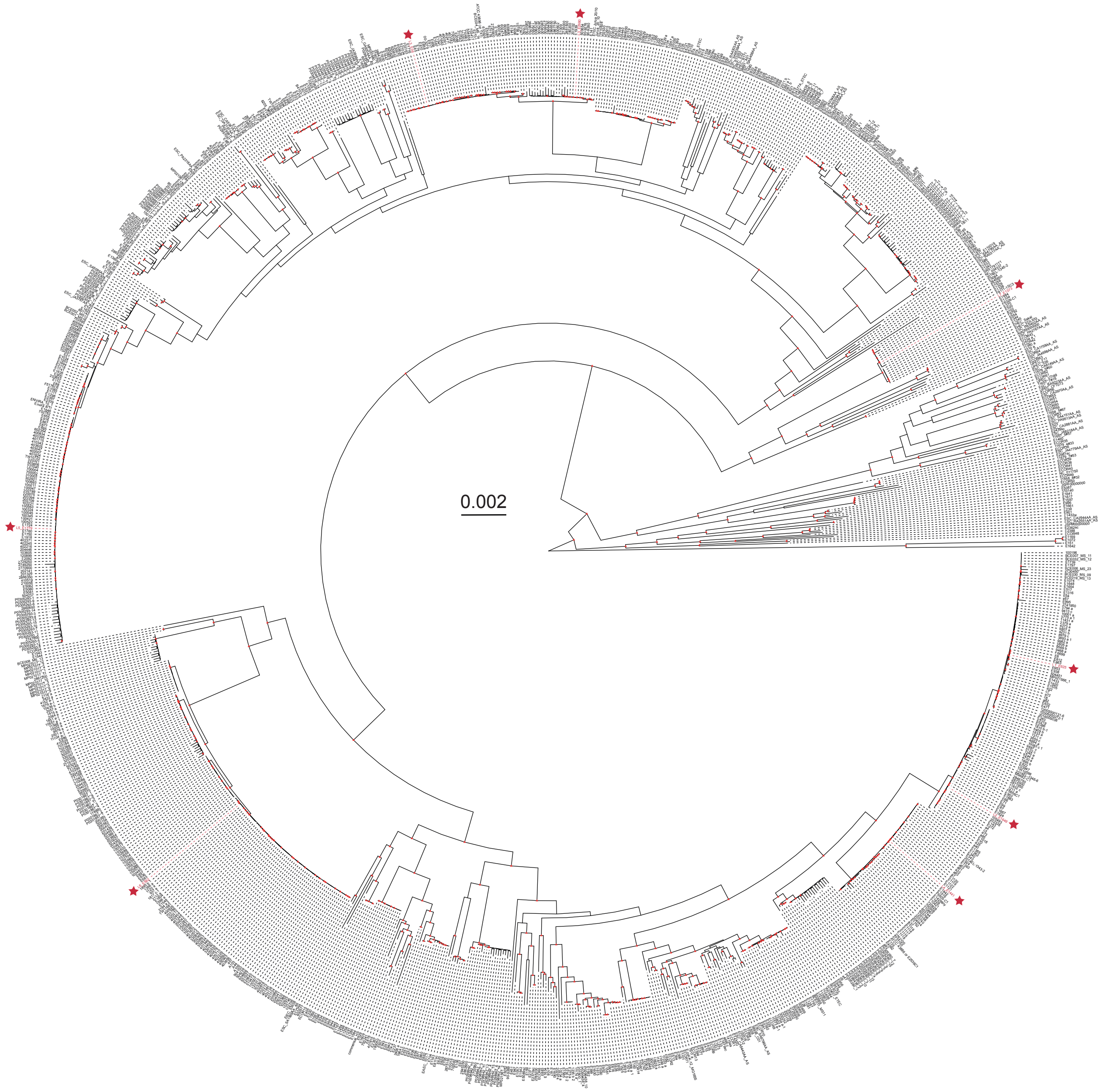


**Figure S11:** Bootstrap values of the phylogenetic tree of ETEC and *E. coli* isolates. Maximum likelihood phylogeny (1000 ultra fast parametric bootstraps, midpoint rooted). Nodes with a bootstrap values >95 are marked with a red triangle. Strain IDs are located on the tips and the ETEC reference strains are colored in red and marked with a red star. The scale bar indicates the number of substitutions per site.

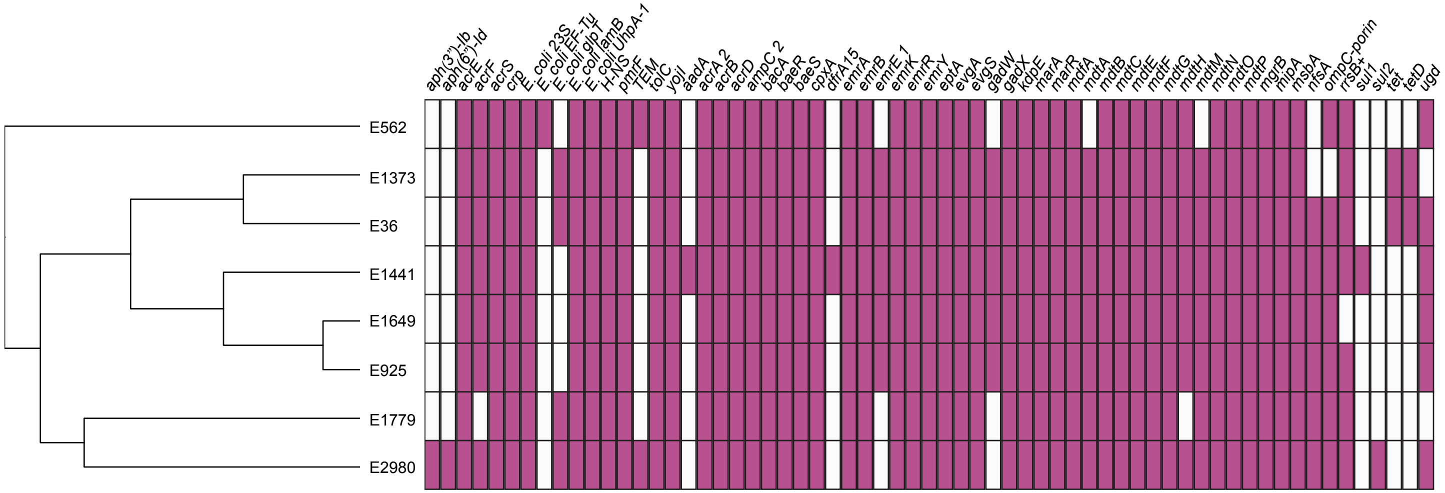


**Figure S12:** Presence and absence of antibiotic resistance genes.

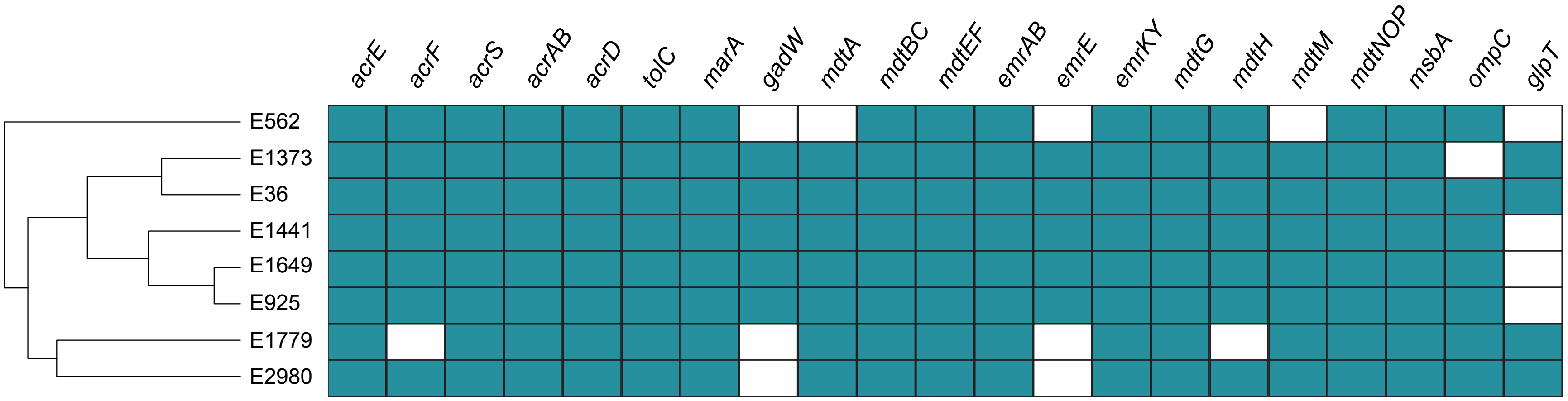


**Figure S13:** Presence and absence of efflux systems and porins.

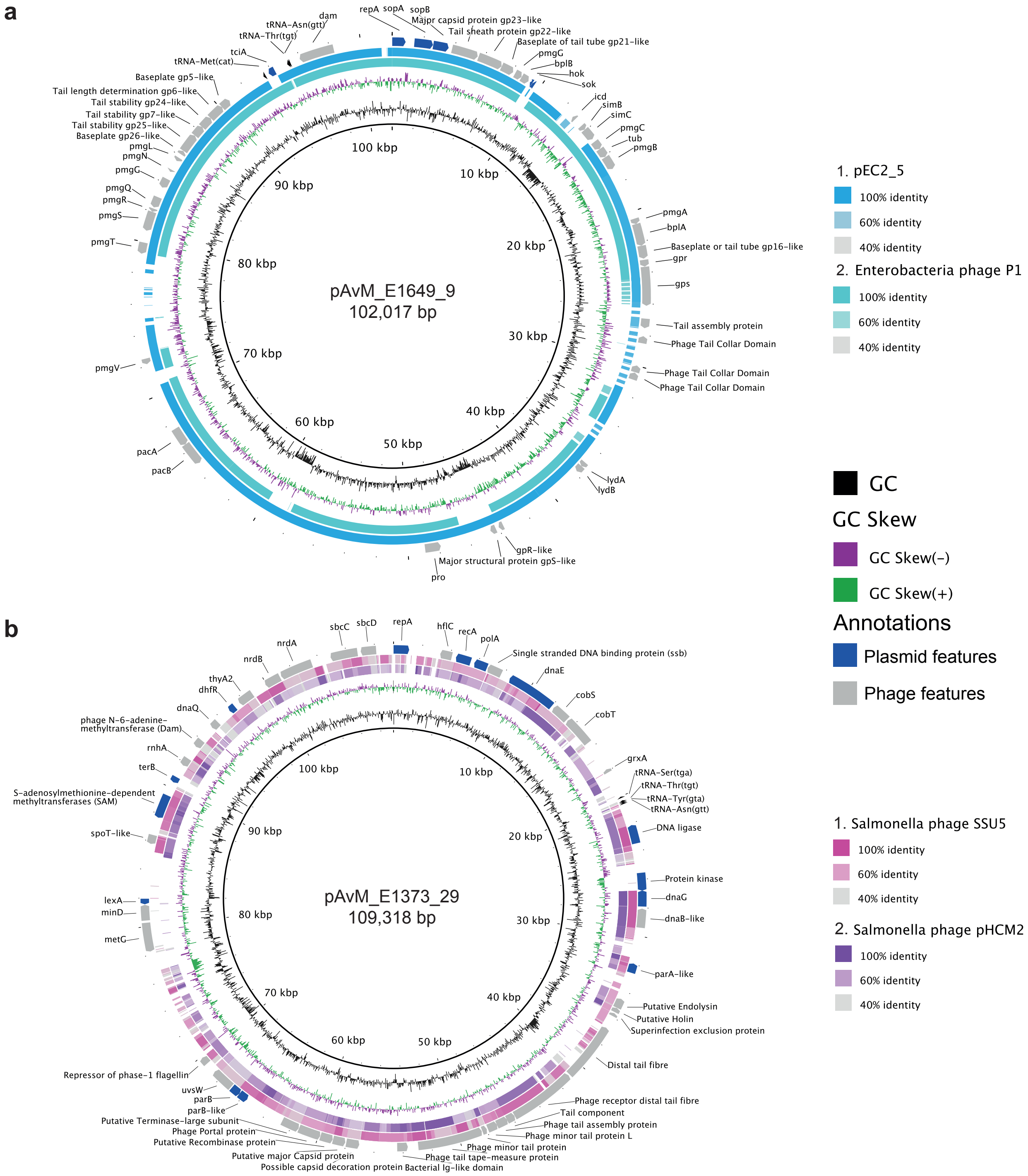


**Figure S14:** Comparison between the two identified ETEC phage-plasmids using tblastx. **a)** pAvM_E1649_9 is a P1-like phage plasmid here compared to pEC2_5 (*E. coli* strain EC2_5; CP041960.1) and Enterobacteria phage P1 (*Escherichia* virus P1; NC_005856.1). **b)** pAvM_E1373_29, a phage-plasmid similar to the SSU5 (Salmonella phage; JQ965645.1) and pHCM2 (*Salmonella typhi* strain CT18; AL513384.1) in *Salmonella typhi.* Tblastx comparison were made using BRIGS with the thresholds indicated to the right of each plasmid comparison. Annotations are from the respective ETEC phage-plasmids which were used as references for the blast.

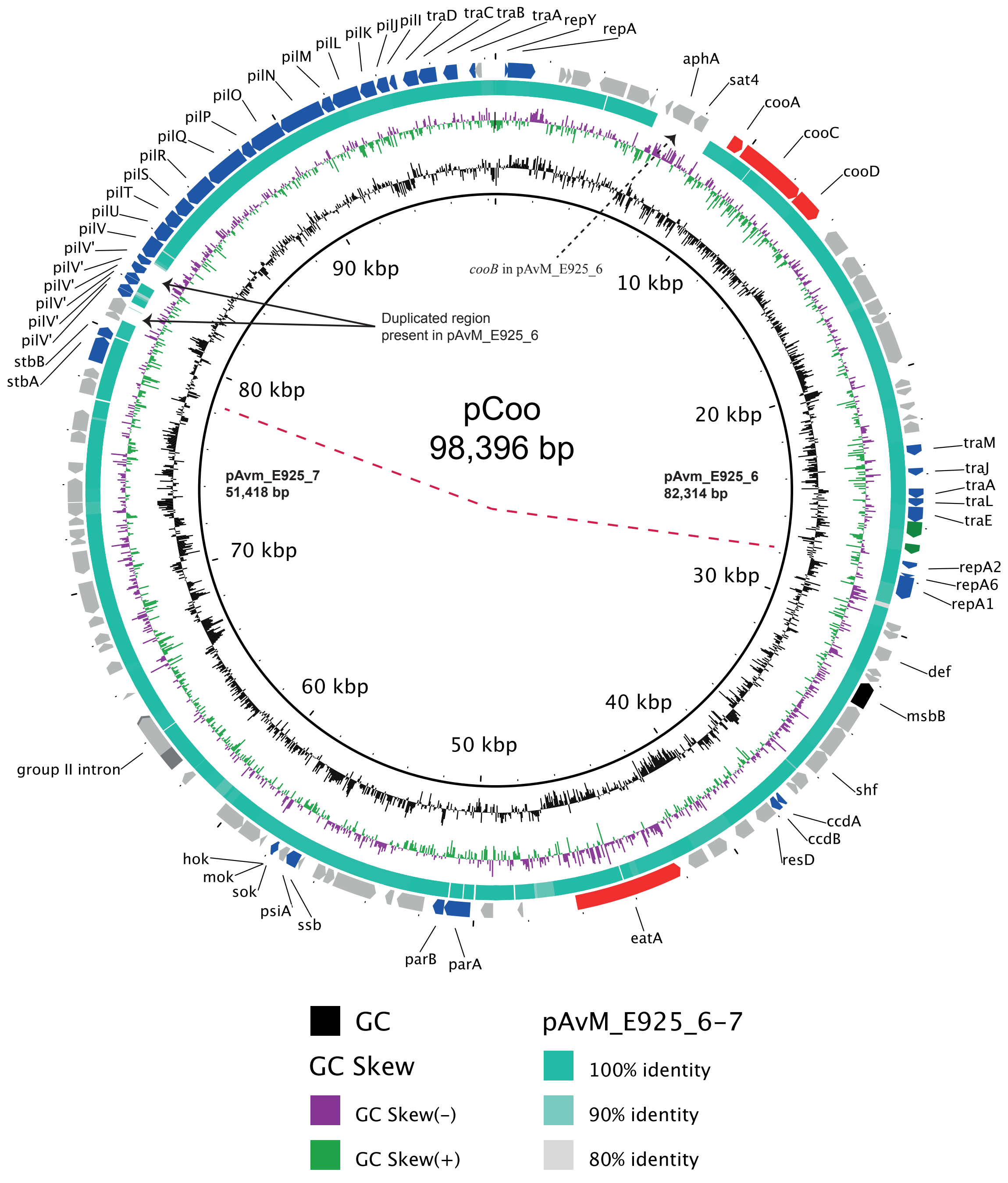


**Figure S15:** BLASTn comparison between the mosaic cointegrate pCoo (AY536429.1) and the concatenated plasmids pAvM_E925_6 and 7. The red dashed line indicates which part of the pCoo matches with which ETEC reference plasmid. An insertion and duplication have been introduced in pAvM_E925_6 as indicated by the two arrows. The gene *cooB* is missing from pCoo but is present in pAvM_E925_6, indicated with a dashed arrow.

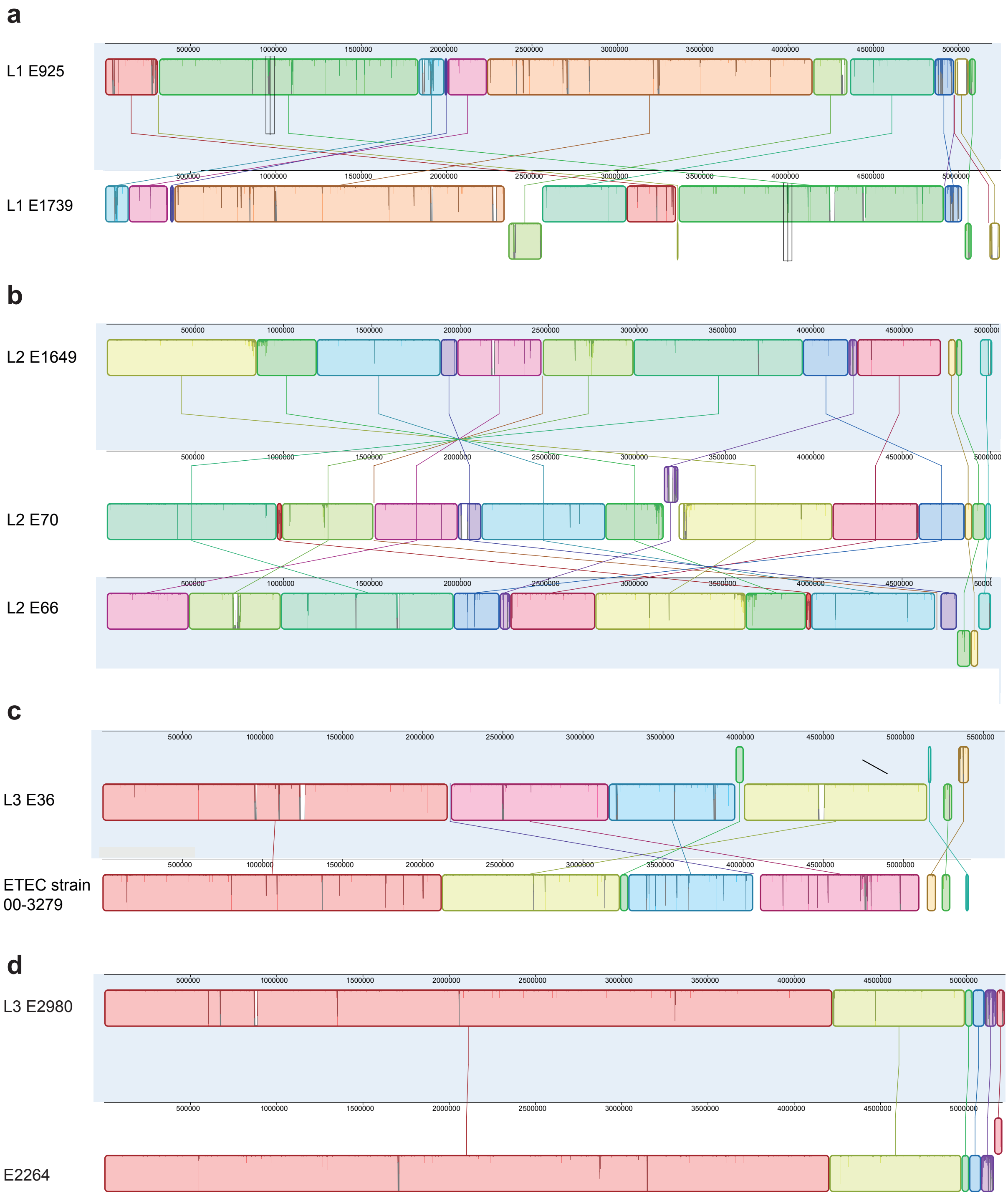
a
L1 E925
L1 E1739
b
L2 E1649
L2 E70
L2 E66
c
L3 E36
ETEC strain 00-3279
d
L3 E2980
E2264

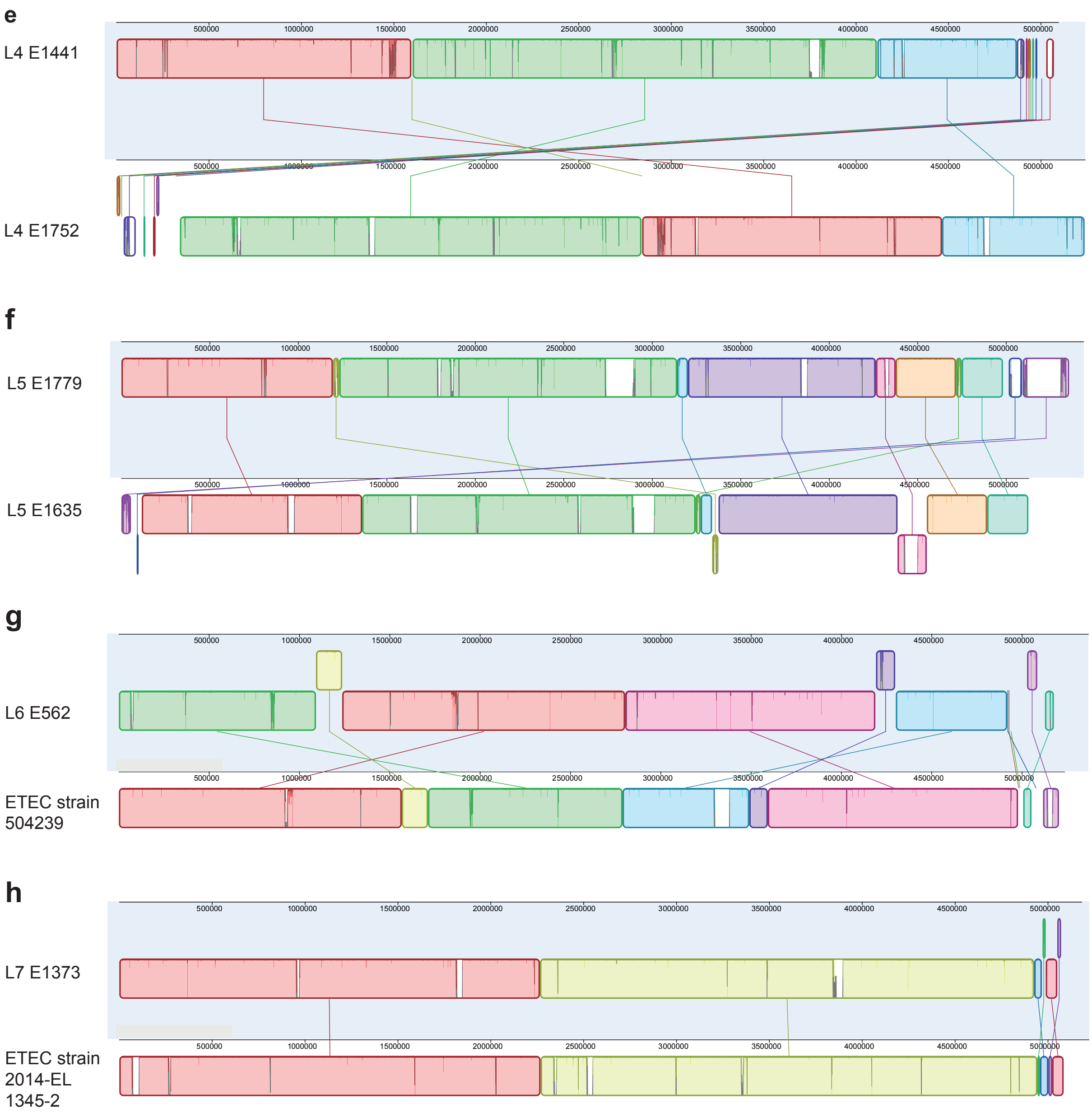


**Figure S16:** Whole genome comparisons using progressiveMAUVE between each ETEC reference strain with one or two different ETEC strains that belong to the same lineage. **a)** ETEC reference strain E925 compared to another long-read sequenced ETEC strain (E1739) that belong to the same lineage (L1). **b)** ETEC reference strain E1649, lineage 2, compared to two additional L2 ETEC strains (E70 and E66). **c)** ETEC reference strain E36, lineage 3, compared to ETEC strain 00-3279 (CP024293-96). **d)** ETEC reference strain E2980, lineage 3, compared to E2264 (CP023349-52). **e)** ETEC reference strain E1441 compared to E1752, both part of lineage L4. **f)** ETEC reference strain E1779 compared to E1635 part of lineage 5. **g)** ETEC reference strain E562 from lineage 6 compared to the ETEC strain 504239 (CP025859-61). **h)** ETEC reference strain E1373 part of lineage 7 compared to the ETEC strain 2014-EL-1345-2 (CP024223-27).

**Table S1:** General characteristics of the chromosome of the ETEC reference strains

| Strain | L1 E925 | L2 E1649 | L3 E36 | L3 E2980 | L4 E1441 | L5 E1779* | L6 E562 | L7 E1373 |
|---|---|---|---|---|---|---|---|---|
| **Length (bp)** | 4,858,376 | 4,721,269 | 5,151,162 | 4,992,286 | 4,869,798 | 4,982,341 | 4,918,084 | 4,926,854 |
| **GC (%)** | 50.7 | 50.8 | 50.7 | 50.7 | 50.8 | 50.8 | 50.9 | 50.4 |
| **Total no of CDSs** | 4,521 | 4,409 | 4,816 | 4,924 | 4,560 | 4,649 | 4,568 | 4,760 |
| **Accession number** | LR883050 | LR882973 | LR882997 | LR882978 | LR883012 | LR883006 | LR883000 | LR882990 |

* Chromosome not circularized

**Table S2:** Phenotypic antibiotic profile of the ETEC reference strains.

| **Strain** | **Lineage** | **MIC mg/L (R/I/S)*** | | | | | | | | | | | | | |
|---|---|---|---|---|---|---|---|---|---|---|---|---|---|---|---|
| | Class | Penicillin | | | Cephalosporin | | Tetracyclines | | Fluoroquinolones | | Macrolides | | Sulphonamide | Misc agents | |
| | Antibiotic | Amp | Amc* | Oxa | Caz | Cro | Dox* | Tet* | Na* | Nor | Azm* | Ery* | Sxt | Cm | NIT |
| **E925** | L1 | 4 (S) | 4 (S) | >128 | 0.5 (S) | 0.5 (S) | 2(S) | 8 (I) | 32 (R) | 0.5 (S) | 16 (S) | 16 | < 0.25 (S) | 8 (S) | 32 (S) |
| **E1649** | L2 | 8 (S) | 8 (S) | >128 | 0.5 (S) | 2 (I) | 2 (S) | 64 (R) | 16 (S) | 2 (S) | 16 (S) | 64 | 0.25 (S) | 16 (I) | 32 (S) |
| **E36** | L3 | 16 (I) | 16 (I) | >128 | 0.5 (S) | 2 (I) | 64 (R) | 2 (S) | 16 (S) | 1 (S) | 16 (S) | 64 | S) | 8 (S) | 16 (S) |
| **E2980** | L3 | >128 (R) | 16(I) | >128 | 0.5 (S) | 2 (I) | 2 (S) | 16 (R) | < 0.25 (S) | 2 (S) | 64 (R) | 128 | 0.25 (S) | 16 (I) | 32 (S) |
| **E1441** | L4 | 4 (S) | 4 (S) | >128 | 0.5 (S) | 2 (I) | 8 (I) | 128 (R) | 8 (S) | 1 (S) | 16 (S) | 64 | >128 (S) | >128 (R) | 16 (S) |
| **E1779** | L5 | 16 (I) | 16 (I) | >128 | 0.5 (S) | 2 (I) | 2 (S) | 2 (S) | 16 (S) | 2 (S) | 16 (S) | 16 | < 0.25 (S) | 16 (I) | 16 (S) |
| **E562** | L6 | >128 (R) | 16(I) | >128 | 0.25 (S) | 1 (S) | 8 (I) | 128 (R) | 8 (S) | 0.5 (S) | 16 (S) | 32 | < 0.25 (S) | 8 (S) | 32 (S) |
| **E1373** | L7 | 8 (S) | 8 (I) | >128 | 0.5 (S) | 2 (I) | 64 (R) | >128 (R) | 8 (S) | 2 (S) | 16 (S) | 64 | < 0.25 (S) | 16 (I) | 32 (S) |

Amp = Ampicillin, Amc = Amoxicillin-clavulanic acid, Oxa = Oxacillin, Caz = Ceftazidime, Cro = Ceftriaxone, Dox = Doxucycline, Tet = Tetracyclin, Na = Naldixic acid, Nor = Norfloxacin, Azm = Azithromycin, Ery = Erythromycin, Cm = Chloramphenicol, NIT = Nitrofurantoin Sxt = sulphamethoxazole-trimethoprim

R = resistant; I = Intermediate; S = sensitive

*Clinical MIC breakpoints were not determined by EUCAST (The European Committee on Antimicrobial Susceptibility Testing. Breakpoint tables for interpretation of MICs and zone diameters. 2016., Version 9. http://www.eucast.org. [Accessed Febrary 2017] and The Clinical & Laboratory Standards Institute (CLSI).

**Table S4:** Identified prophages in the chromosome and plasmids in each of the reference ETEC strains. Region of prophages and potential cargo genes are indicated.

| Lineage/Strain/CF profile | Chromosome/ plasmid | Prophage (Pph) | Source | Location[a] | Size (kB) | GC % | No. of proteins | Cargo genes[b] |
|---|---|---|---|---|---|---|---|---|
| **L1 E925 CS1+CS3+CS21** | Chromosome | E925_Pph_1 | Entero_P88_ NC_026014(25) | 1980800-2010357 | 29.6 | 50.1 | 39 | flagella biosynthesis regulator (E925_01987) |
| | | E925_Pph_2 (Cryptic) | Entero_lambda_ NC_001416(6) | 2829055-2849450 | 23.4 | 50.4 | 24 | *ompX* and *sohB* (probable Protease) |
| | | E925_Pph_3 (cryptic) | Entero_lambda_ NC_001416(4) | 3229100-3240718 | 11.62 | 44.1 | 15 | Outer membrane lipoprotein, *blc* |
| | | E925_Pph_4 | Shigel_SfIV_ NC_022749(12) | 3724871-3749140 | 24.3 | 46.7 | 32 | *gadW* transcriptional regulator of the AraC family (helix-turn-helix motif). tRNA-Thr insertion site |
| | pAvM_E925_7 | E925_Pph_5 | Stx2_c_1717_ NC_011357(3) | 23316-45489 | 22.1 | 48.7 | 22 | *eatA_3, eatA_4, eatA_5* (*eatA* disrupted in three ORFs), *stbAB* and *psiA* |
| **L2 E1649 CS2+CS3+CS21** | Chromosome | E1649_Pph_1 Cryptic phage | Escher_TL_2011b NC_019445(29) | 1298543-1335201 | 36.6 | 49.1 | 45 | *istB*-like ATP binding protein similar to DNA replication protein, *dnaA*. |
| | | E1649_Pph_2 | Entero_P88_ NC_026014(33) | 1502430-1542242 | 29.1 | 52.4 | 46 | Putative Acetyltransferase and arabinose transporter |
| | | E1649_Pph_3 (cryptic phage) | Entero_P88_ NC_026014(15) | 2177200-2195400 | 17.7 | 49.9 | 23 | No cargo genes |

| | | | | | | | | |
|---|---|---|---|---|---|---|---|---|
| | | E1649_Pph_4 | Entero_P4_ NC_001609(9) | 4048900-4077099 | 21.0 5 | 48.4 | 26 | *nanC*- Porin gene (N-acetylneuraminate epimerase precursor and putative inner membrane protein). tRNA-Leu insertion site |
| | pAvM_E1649_ 9 | E1649_Pph_6 | Salmon_SJ46_ NC_031129(8 8) | 4916371-5038771 | 122. 4 | 47 | 138 | *eatA*. This is the P1 like phage-plasmid |
| **L3 E36 CFA/I + CS21** | Chromosome | E36_Pph_1 (cryptic phage) | Entero_P4_ NC_001609(9) | 47658-59450 | 11.6 | 48 | 15 | Retron-type Reverse transcriptase and DNA primase |
| | | E36_Pph_2 | Entero_P4_ NC_001609(9) and Salmon_Fels_ 2 NC_010463(3 7) | Two adjacent phages: 1220220-1231640 and 1231677-1266590 | 46.4 | 49.6 | 61 | Putative bacteriocin adjacent to the *intA_2* gene (E36_01826). Insertion site is in the region of *ssrA* |
| | | E36_Pph_3 | Entero_P88_ NC_026014(4 1) P2 family of bacteriophage | 1285100-1328400 | 43.3 | 52.7 | 56 | stbA and stbB (plasmid stability genes) and *tfp* pilus assembly gene*, pilA* |
| | | E36_Pph_4 | Entero_mEp46 0_ NC_019716(2 3 | 2138535-2199525 | 48.0 | 50.3 | 63 | *terB* (tellurite reistance), and *ompX.* Insert site is tRNA-Gly |
| | | E36_Pph_5 | Entero_mEp46 0_ NC_019716(2 3) (probable cryptic phage) | 2499123-2535609 | 36.5 | 50.9 | 40 | No cargo genes |

| | | | | | | | | |
|---|---|---|---|---|---|---|---|---|
| | | E36_Pph_6 | Burkho_Bcep Mu_NC_0058 82(32) | 2738731-2795496 | 56.7 | 52.4 | 68 | Potassium transporter, *trkG* and *parB* partition protein (with nuclease N-terminal domain) |
| | | E36_Pph_7 | Entero_HK62 9_ NC_019711(2 1) | 3024129-3076300 | 36.5 | 51.3 | 56 | Tellurite resistance protein and ncRNA, *dicF* and *dicC* genes present are of interest but may not be considered as cargo). |
| | | E36_Pph_8 | Entero_lambd a_ NC_001416(2 7) | 3532592-3576100 | 43.5 | 51.6 | 71 | Outer membrane protein Blc. Group II intron reverse transcriptase (*ltrA*_14) with adjacent group II intron *uvrB*, *sohB_3, dicC* (E36_04216) |
| | | E36_Pph_9 | Entero_P4_ NC_001609(1 2) | 44922786-4519904 | 27.6 | 49.8 | 14 | No cargo genes |
| | pAvM_E36_12 | E36_Pph_10 (cryptic phage) | Entero_BP_47 95_ NC_004813(3) | 182814-195964 | 13.1 | 52.9 | 23 | No cargo genes |
| | | E36_Pph_11 (cryptic phage) | Entero_BP_47 95_ NC_004813(3) | 267737-281095 | 13.3 | 51.2 | 21 | *sopB* and *estA2* |
| | Prophage* Contig 2 | E36_Pph_12 (Partial - no *int* gene present. Probable cryptic) | Salmon_Fels_ 2_ NC_010463(2 4) | 700-18544 | 17.8 | 55.4 | 29 | No cargo genes |
| | Prophage* Contig 998 | E36_Pph_13 Cryptic phage) | Entero_lambd a_ NC_001416(1 1 | 1-16565 | 16.5 | 54.2 | 22 | No cargo genes |

| | | | | | | | | |
|---|---|---|---|---|---|---|---|---|
| **L3 E2980 CS7** | Chromosome | E2980_Pph_1 | Salmon_SEN34_NC_028699(24) | 1314690-1354910 | 40.22 | 49.7 | 50 | No cargo genes |
| | Chromosome | E2980_Pph_2 | Shigel_Sf6_NC_005344(14) | 1568700-1611651 | 42.9 | 46.4 | 63 | *wzy* (E2980_01970) gene is an oligosaccharide repeat unit polymerase adjacent to the integrase (*oac* gene in sf6 phage at the same location next to *int*).tRNA-Arg insertion site |
| | Chromosome | E2980_Pph_3 | Entero_P88_NC_026014(43) | 2070150-2109664 | 39.5 | 51.6 | 55 | Putative plasmid stability proteins and *repA* (E2980_02444, E2980_02445 and E2980_02446) and flagella biosynthesis regulator (as found in the P88 phage of E925 above) near *int* gene. tRNA-Leu insertion. This pahge is related to P1 phage so it can probably exist as a Phage-Plasmid. |
| | Chromosome | E2980_Pph_4 (Cryptic phage) | Entero_lambda_NC_001416(11) | 2447775-2466400 | 18.6 | 51.8 | 17 | No cargo genes |
| | Chromosome | E2980_Pph_5 | Entero_lambda_NC_001416(22) | 2918700-2962650 | 43.95 | 51.6 | 60 | *tciAB* (tellurite resistance), *ompX* |
| | Chromosome | E2980_Pph_6 | Entero_lambda_NC_001416(27) | 3588788-3636557 | 47.8 | 49.6 | 58 | No cargo genes. tRNA insertion site adjacent to *int* gene, *ompX* gene |

| | | | | | | | | |
|---|---|---|---|---|---|---|---|---|
| | Prophage* Contig 5 | E2980_Pph_7 | Salmon_SEN34_ NC_028699(13) | 8730-21920 | 13.1 | 49.4 | 25 | No cargo genes |
| **L4 E1441 CS6** | Chromosome | E1441_Pph_1 | Salmon_HK620_ NC_002730(17) | 1473781-1517255 | 43.5 | 46.5 | 57 | tRNA-Arg insertion site. CatB-related O-acetyltransferase adjacent to *int* |
| | Chromosome | E1441_Pph_2 | Entero_mEp460_ NC_019716(11) | 2663330-2710354 | 47.0 | 50.6 | 65 | Anti-adapter gene *iraM* (operates via *rpoS*) |
| | Chromosome | E1441_Pph_3 | Entero_lambda_ NC_001416(26) | 3194858-3250061 | 55.2 | 50.4 | 66 | Phospholipid binding protein, *ybhB*; Inner membrane protein, yeaI; *slyB* lipoprotein precursor, *sohB;* outer membrane protein, *ompD and slyB* and *kilR* |
| | Chromosome | E1441_Pph_4 | Entero_mEp460_ NC_019716(37) | 3756217-3834601 | 78.38 | 52 | 101 | CysO- .Phage adjacent to tRNA-Thr |
| **L5 E1779 CS5+CS6** | Chromosome | E1779_Pph_1 | Salmon_Fels_2_ NC_010463(36) | 1195952-1232915 | 36.9 | 49.1 | 45 | Cargo genes are present but all the hypothetical. *ssrA* integration site. |
| | Chromosome | E1779_Pph_2 | Entero_P2_ NC_001895(21) | 1783098-1807397 | 24.3 | 49.8 | 32 | No cargo genes identified |
| | Chromosome | E1779_Pph_3 | Entero_lambda_ NC_001416(22) | 2368430-2397000 | 28.57 | 55.4 | 37 | *sohB*-probable protease |

| | | | | | | | | |
|---|---|---|---|---|---|---|---|---|
| | Chromosome | E1779_Pph_4 | Entero_lambda_ NC_001416(18) | 3157352-3201305 | 43.9 | 51.6 | 56 | *iraM* (anti-adaptor protein blocking RpoS degradation) |
| | Chromosome | E1779_Pph_5 | Entero_cdtI_NC_009514(22) | 2859100-2906257 | 47.16 | 51.6 | 65 | Some potential cargo genes but all hypothetical |
| | Chromosome | E1779_Pph_6 | Entero_lambda NC_001416(29) | 3288450-3335837 | 47.4 | 51.5 | 65 | Putative *ompD* porin protein, E1779_03897 encodes ZapA, a cell division protein |
| | Chromosome | E1779_Pph_7 | Shigel_SfII NC_021857(35) | 3839659-3878224 | 38.5 | 50.8 | 55 | tRNA-Thr insertion site. Region with number of regulators: HTH-type transcriptional regulator *gadW*-like (acid resistance).lexA repressor, *perC* activator, *gntR* family regulators, *yfdR* family of *dicC* DNA binding transciprtional regulator, lexA/HTH-type transcriptional regulator; *yfbR*- 5' deoxynucleotidase |
| | Chromosome | E1779_Pph_8 | Entero_fiAA91_ss_NC_022750(31) | 4718700-4750300 | 31.6 | 51.7 | 42 | No cargo genes |
| | pAvM_E1779_19 | Phage islet | Stx2_c_1717_ NC_011357(3) | 49007-62586 | 13.5 | 51.3 | 23 | *estA3/4*, *traI*, *traX*, *finO* |
| | Probable phage-plasmid | E1779_Pph_9 | Entero_N15_ NC_001901(21) | 1-46243 | 46.2 | 50.5 | 69 | *ompX* (resistance to complement killing), Phage-specific genes |

| | | | | | | | | |
|---|---|---|---|---|---|---|---|---|
| **L6 E562 CFA/I+CS21** | Chromosome | E562_Pph_1 | Entero_lambda_ NC_001416(22) | 2695572-2745700 | 49.4 | 51.5 | 62 | *ftsZ* inhibitor gene, *kilR*. Though may be of phage origin. |
| | Chromosome | E562_Pph_2 | Entero_fiAA91_ss NC_022750(31) | 4666803-4698527 | 31.7 | 52 | 42 | No cargo genes |
| **L7 E1373 CS6** | Chromosome | E1373_Pph_1 | Entero_mEp460 NC_019716(23) | 2848300-2890275 | 41.98 | 51.5 | 22 | *ompX* (resistance to complement killing), Tellurite resistance gene, *tehB and a S*-Adenosyl-l-methionine (SAM)-dependent |
| | Chromosome | E1373_Pph_2 | Shigel_SfIV NC_022749(34) Similar to the content of E1779_Pph_7 (Shigel_SfII_ NC_021857(35)) | 3856500-3897232 | 40.73 | 48.7 | 59 | tRNA-Thr insertion site. *kilA*-like domain, *lexA, perC* and *gntR* regulator. An almost identical region is present in the *Shigella* phage related to SfIV (accession number KC814930.1) |
| | pAvM_E1373_29 | E1373_Pph_3 | Salmon_SSU5_ NC_018843(91) This is the Phage-Plasmid related to pHCM2 and SSU5 | 1-109318 | 109.1 | 46.4 | 130 | *terB*, mind (cell division inhibitor), *uvrB*, *parB*-like (2 genes, ORFs:04807 and 04808), *hlyB*, Flp pilus assembly (ORF: 04842), *grxA* (glutaredoxin), *cobT* and *cobS* (aerobic cobaltochelatase subunits), *dnaE, polA* (DNA |

| | | | | | | | | |
|---|---|---|---|---|---|---|---|---|
| | | | | | | | | polymerase I), recA (recombiase A), *repA* (FIB), *nrdA* and *nrdB* (ribonucleoside-diphoasphatase), a lot of hypo proteins. |

[a]The location within the chromosome, plasmid or contig individually.

[b]The locus_tags will be updated when sequences have been accepted in GenBank.

**Table S3:** Genomic antibiotic resistance profile, including plasmid and chromosomal genes as well as efflux systems and porins and metal resistance

| **Strain** | **Lineage** | **Antibiotic resistance genes** | **Porins and efflux pumps associated with antibiotic resistance*** | **Metal resistance** |
|---|---|---|---|---|
| **E925** | L1 | *ampC*, ampDE** | *acrRAB-tolC, acrEF-S, crp, emrRAB, emrE, mdf(A), gadW, mdtAB, mdtEF, mdtG, mdtH, mdtM, mdtNOP, msbA, ompC* | |
| **E1649** | L2 | *ampC*, ampDE** | *acrRAB-tolC, acrEF-S, crp, emrRAB, emrE, gadW, mdf(A), mdtAB, mdtEF, mdtG, mdtH, mdtM, mdtNOP, msbA, ompC* | |
| **E36** | L3 | *ampC*, ampDE*, tet(B)* | *acrRAB-tolC, acrEF-S, crp, emrRAB, emrE, gadW, glpT, mdf(A)-like, mdtAB, mdtEF, mdtG, mdtH,mdtM, mdtNOP, msbA, ompC* | |
| **E2980** | L3 | *ampC*, ampDE*, blaTEM-1b, strA (aph(3'')-Ib-like), strB (aph(6)-Id), sul2* | *acrRAB-tolC, acrEF-S, crp, emrRAB, glpT, mdf(A), mdtAB, mdtEF , mdtG, mdtH, mdtM, mdtNOP, msbA, ompC* | |
| **E1441** | L4 | *aadA1-like, ampC*, ampDE*, dfrA15, sul1 (Class I integron), tet(A) (Tn1721)* | *acrRAB-tolC, acrEF-S, crp, emrRAB, emrE, gadW, mdf(A)-like, mdtAB, mdtEF, mdtG, mdtH, mdtM, mdtNOP, msbA, ompC* | *mer* operon (*merRTPCADE*) (Tn21), *qacE* (part of the Class I integron) |
| **E1779** | L5 | *ampC*, ampDE** | *acrRAB-tolC, crp, emrRAB, glpT, mdf(A)-like, mdtAB, mdtEF, mdtG, mdtM, mdtNOP, msbA* | |
| **E562** | L6 | *ampC*, ampDE*, blaTEM-1b, tet(A)* | *acrRAB-tolC, acrEF-S, crp, emrRAB, mdf(A)-like, mdtEF, mdtG, mdtH, mdtNOP, msbA, ompC* | *mer* operon (*merRTPCADE*) (Tn21) |
| **E1373** | L7 | *ampC*, ampDE*, tet(B)* | *acrRAB-tolC, acrEF-S, crp, emrRAB, emrE, glpT, mdf(A)-like, mdtAB, mdtEF, mdtG, mdtH, mdtM, mdtNOP, msbA* | |

* Located on the chromosome
