## Additional file 2 for "Long-read-sequenced reference genomes of the seven major lineages of enterotoxigenic *Escherichia coli* (ETEC) circulating in modern time"

### Additional file 2
### Detailed description of ETEC reference plasmids

**ETEC strain E925 (L1)**
The reference strain E925 expresses the CFs CS1, CS3 and CS21 along with the enterotoxins LT and STh and belong to lineage L1 [1]. Members of L1 have the same virulence profile. Genomic analysis revealed 4 plasmids in total, pAvM_E925_4-7. Interestingly, two or possibly three of the identified plasmids (pAvM_E925_4, 6 and 7) may be a co-integrated plasmid where individual replicons are present (8). Several plasmids were assigned to the same replicon type, IncFII (pAvM_E925_4, pAvM_E925_5 and pAvM_E925_7). The ETEC plasmid pCoo is a co-integrate with two replicons, IncFII and IncI1. A blastn comparison of the concatenated pAvM_E925_6 and pAvM_E925_7 show that these most likely form a co-integrate plasmid similar to pCoo (Figure S15).

**pAvM_E925_4**
The pAvM_E925_4 is predicted as an IncFII (FII-11) plasmid with a predicted *oriT* site. The prediction is supported by its size (116,803b) and because of an intact conjugation machinery (*tra* genes) which is similar to R100 (the quintessential IncFII plasmid) [2]. Blastn analysis revealed that the plasmid is similar to a number of plasmids of sequenced O6:H16 ETEC isolates including the CS3+ plasmid in the reference strain E1649 (Figure 1a). Specifically, plasmid unnamed2 of PacBio sequenced ETEC isolate F5656C1 isolated by CDC in US [3], a plasmid of ETEC isolate FMU073332 isolated in Mexico 1987 (CP017848.1) and plasmid pETEC_80 of type strain E24377A.

**Plasmid features:** A partial *tra* region is present, but the plasmid may be mobilizable with the help from other plasmids. The plasmid contains genes involved in plasmid stability (*psiAB*, *stbAB*). Two TA systems are present in this plasmid: *ccdAB* system, where *ccdA* encoding the toxin and *ccdB* which encodes the antitoxin needed for survival of the cell, and the *hok-sok* plasmid antisense RNA-regulated system.
**Virulence:** The pAvM_E925_4 harbours the gene cluster encoding CS3 (*cst* genes) as well as *eltAB* (LTh variant LT1) and STh (e*stA3/4)*. The CS3 locus consist of 8 CDS, *cstA-H*. The *cstA* encodes a chaperone and *cstG* and *cstH* encodes two major subunits. The outer membrane usher is encoded by *cstB* which also contain CDSs for four additional proteins: *cstC-F*. The C-terminal of *cstB* is missing compared to the reference [4] and is found in the second open reading frame. A nucleotide comparison between the *cstB* in E925 and *cstB* of additional CS3 positive strains (CS1+CS3+/-CS21, CS2+CS3+/-CS21) [1] as well as *cstB* in X16944.1, to the available reference sequence encoding *cstC* (63 kD protein found within *cstB*) show that an extra G is inserted at position 2432 in *cstB* of all analysed sequences. This generates a frame-shift mutation resulting in a premature stop codon and the C-terminal to be found in the second reading frame instead of the first. In addition to the classical ETEC virulence genes this plasmid also contain the genes encoding for the putative virulence factor EtpBAC [5–7]. The gene encoding the ABC transcriptional regulator Rns is located on this plasmid and required for expression of CS1 or CS2 colonization factors and for adhesion [8].

**pAvM_E925_5**

**Inc group:** This is a multi-replicon plasmid with both IncFII (FII-111) and IncFIB (FIB-45). Blastn analysis showed that pAvM_E925_5 is most similar to a plasmid (CP024258.1) from the ETEC strain F5505-C1 [3] with 80% query coverage and 98.9% sequence identity as well as the p557 of the CS1+CS3+CS21 positive reference strain E1392/75 (FN822746) (12) with 68% coverage with 99.9% sequence identity. The p557 does not contain the conjugal machinery which pAvM_E925_5 does. The conjugal transfer genes were found to be 99% identical to the corresponding genes in the multi-drug resistant pHK17a-like plasmid present in an *E. coli* strain isolated from a patient with a bloodstream infection, specifically a ST95 *E. coli*, which is a multi-drug resistant plasmid (JF779678.1).

**Plasmid features:** Harbours an intact *tra* region as a well as a predicted *oriT*. Genes involved in plasmid stability are present (*psiAB, parB* and *sopAB*).

**Virulence:** The locus (*lngX1, R, S, T, X2, A-J, P*) encoding the colonisation factor CS21, one of the most prevalent CFs, is present in this plasmid.

**pAvM_E925_6**

The plasmid was assigned to IncI1 and may be part of a co-integrated plasmid, most likely together with pAvM_E925_7 which is assigned to IncFII (FII-15) (Figure S6). Blastn comparison showed highest homology to p746 from ETEC strain E1392/75 (FN822748.1) with 96% query coverage and 99.9% nucleotide identity and share sequence homology to pCoo (CR942285) from the ETEC strain JEF100 showed an 80% coverage at a >99% identity. In pAvM_E925_6 there is a duplicated region a region encompassing a shufflon-specific DNA recombinase (*rci*), *traE-G*, *traM*, *nikB* and *nikA* have been duplicated and upon this duplication the *traE* was interrupted due to a deletion. The same region is present in p746 of 1392/75 but not present in the pCoo plasmid. The duplication has resulted in the presence of two *oriT* regions (Figure 1b). Nucleotide alignment show that large regions are shared across pAvM_E925_6 and pCoo, however the pAvM_E925_6 lacks part of the *tra* locus (blastn comparison with the IncI1 R64 plasmid). Furthermore, the pAvM_E925_6 plasmid contains an intact *pil* loci encoding for the thin pilus and is 99% identical to the *pil* loci in R64.

**Plasmid features:** Genes involved in plasmid stability were identified; *stbAB* as well as the *vapBC* toxin-antitoxin system. The plasmid harbours the *iib* gene encoding the Colicin 1b immunity protein, however it lacks the additional structural *cib* gene.

**Virulence:** The *coo* loci (*cooABCD*), which encodes the proteins needed for CS1, is located on this plasmid. The putative virulence factor *cexE* and the *aat* operon is also present.

**pAvM_E925_7**

This plasmid was assigned to IncFII (FII-15) and share part of the CS1 plasmid pCoo (Figure S6).

**Plasmid features:** The plasmid only carries five *tra* genes (*traM*, *traA*, *traL*, *traE* and *traK*). This may be an indication that this plasmid is part of a co-integrated plasmid together with pAvM_E925_6 (IncI1) as described above (see pAvM_E025_6). The following plasmid stability genes are present in the plasmid: *ccdAB*, *psiA*, *repAB* and *stbAB*.

**Phage features:** A complete prophage (E925_Pph_5: Stx2_c_1717_NC_011357) was identified in this plasmid and contains *eatA*_3-5 as cargo genes as well as plasmid stability genes *stbAB* and *psiA* along with multiple IS elements.
**Virulence:** Two CDSs were identified as *eatA_1* and *eatA_2*, sequence alignment showed that this a disrupted *eatA* gene. Downstream of *eatA_1* and *eatA_2* are three additional CDSs predicted as *eatA* (*eatA_3*, *eatA_4*, *eatA_5*), also disrupted by premature stop codons due to two different single base deletions. The multiple *eatA* CDSs are due to a duplication event, including *ygjH* (cyclic di-GMP phosphodiesterase), *eatA* CDSs, ISEc63 and deltaIS*1X2*.

**ETEC strain E1649 (L2)**
The reference strain E1649 expresses CS2, CS3 and CS21. The colonisation factor CS2 is located on the chromosome, whereas CS3 and CS21 are plasmid-borne. Two plasmids were fully circularized: pAvM_E1649_8 and pAvM_E1649_10. A phage-like plasmid was identified but not fully circularised, pAvM_E1649_9. A cryptic plasmid (pAvM_E1649_11) was found but the replicon could not be determined. Four prophages were identified in the chromosome, one of them with the *fim* operon as cargo (Table S5).

**pAvM_E1649_8**
The plasmid is a classical IncFII (FII-15) replicon, however it does not contain a complete *tra* region but an *oriT* was identified. The plasmid is similar to the other CS3+ ETEC reference plasmid pAvM_E925_4 (Figure 1A), although an extra region containing *eatA* is present in pAvM_E1649_8. Furthermore, a region has been duplicated, where in the first region one of the genes encoding MsbB, involved in lipid A biosynthesis, has multiple mutations resulting in premature stop codons. The second region contains the intact *msbB* gene. An *msbB* gene is also located in the chromosome and the plasmid encodes a homologous *msbB* gene. The same as been reported in *E. coli* O157:H7 [9]. Other genes within the duplicated region are the ccdAB (toxin-antitoxin system; plasmid stability), a gene encoding a polysaccharide deacetylase and three hypothetical proteins. The plasmid is similar to pAvM_E925_4, and a 137,665 bp plasmid pEcoFMU07332d (, CP024277.1) isolated from an ETEC strain collected in Mexico 1987 (87% coverage, 99.88% identity). Several other plasmids from O6:H16 strains share similarity with this plasmid (Figure 1a).
Plasmid features: A partial tra region is present as well as the plasmid stability genes *psiAB* and *stbAB*. The plasmid contains the *hok-sok* plasmid antisense RNA-regulated system as well as the *ccdAB* system involved in plasmid stability. Both *hok* (host killing) and *sok* (suppressor of killing) have been identified as having possible roles in selective phage exclusion as well as playing an important role in plasmid stability and maintenance [10].
**Virulence:** The plasmid harbours the CS3 locus, *cstA-H* and the *rns* regulator. Together with the CF, the plasmid also harbours the *eltAB* genes, encoding the LT1 variant (LTh), as well as the *sta3/*4 (STh). The *etpBAC* locus is also located on this plasmid. However, the *etpA* is truncated but may still be functional. Another putative virulence factor is *eatA* which is also present. The plasmid contains the gene for *cexE* located upstream of the *aat* operon.

**pAvM_E1649_9**

This is a P1 phage-like plasmid assigned to IncY that contains the complete prophage E1649_Pph_6 (PHAGE_Salmon_SJ46_NC_031129). pAvM_E1649_9 encodes a full repertoire of genes related to the P1-like phage-plasmid including not only replication protein, *repA* and the partition genes *parA/parB,* but also structural genes for the phage tail, major capsid protein, portal protein, a P1-like toxin/anti-toxin system and various tail fibre proteins. The *tciA* gene encoding a tellurite resistance protein is present, however lacking the full *tciABC* locus which present in the P1 phage. Furthermore *dam*, a methylase, is located upstream of *tciA*. (Figure 2a and Figure S3a). The plasmid is similar to other *E. coli* plasmids; however, these have been isolated from chicken, human blood and urine and no apparent virulence genes could be identified in the plasmids. This suggests that this plasmid has been introduced from a diverse *E. coli* from a different source.
**Plasmid features:** The plasmid stability genes *sopAB* are present.
**Phage features:** A full complement of genes highly similar to P1 bacteriophage.

**pAvM_E1649_10**
The pAvM_E1649_10 plasmid contains two replicons: IncFII (FII-6) and IncFIB (FIB-45) and share sequence homology with pETEC_74 of the ETEC strain E24337A (CP000799.1) (70% query coverage; 99.9% sequence identity) which is a one-replicon plasmid. pAvM_E1649_10 is also highly related to an unknown plasmid from the ETEC strain 2011EL-1370-2 (CP022913.1) which is a two-replicon plasmid (FII and FIB) like pAvM_E1649_10.
**Plasmid features:** A partial *tra* region, plasmid stability genes *psiAB* as well as the plasmid antisense RNA-regulated system *hok-sok* are present.
**Virulence:** The *lng* locus (*lngX1, R, S, T, X2, A-J, P),* encoding the CF CS21, is located on this plasmid.

**pAvM_E1649_11**
A small cryptic plasmid of 8,834 bp harbouring regions similar to a small plasmid in *Klebsiella variicola* strain WCHKP19 plasmid p4_020019 (CP028552). Contains hypothetical genes and two RNA one modulator proteins Rop. No replicons could be identified.

**ETEC strain E36 (L3)**
The strain E36 is one of the two ETEC strains expressing CFA/I. The phylogenetic tree of the ETEC population presented by von Mentzer *et al.*, revealed that there are at least three lineages (L3, L6 and L15) comprising ETEC strains expressing CFA/I [1]. A recent report identified the same lineages but also additional ETEC lineages encompassing CFA/I ST-only strains [11]. In the ETEC strain E36 two plasmids were identified, the largest identified amongst these strains (pAvM_E36_12) categorized as a virulence plasmid, and the smaller plasmid (pAvM_E36_13) categorised as an antibiotic resistance plasmid.

**pAvM_E36_12**
The plasmid is the largest plasmid identified among the eight ETEC reference genomes, a total of 381,858 bp. The size strongly indicates that this a conjugative plasmid as well as the presence of two replicons; IncFII (FII-51) and IncFIB (FIB-10) and potentially a third replicon FIC (FIC-5, novel allele). Due to the size of the plasmid and its content it may be a hybrid

plasmid. The plasmids present in the O78 ETEC strain sequenced by Smith *et al.* [3] show similarity across pAvM_E36_12 (Figure 1c).

**Plasmid features:** This plasmid contains three *tra* regions, some which are complete and others only partial, and multiple copies of several stability genes are present: *psiAB* (6 copies), *repAB* (3 copies), *sopAB* (2 copies) and *stbAB* (4 copies). The plasmid also contains the *hok* and *mok* genes, but lack *sok*, which is a short ssRNA antitoxin.

**Virulence:** The pAvM_E36_12 harbours the CFA/I gene cluster *cfaA*, *B*, *C*, *D* and the CS21 gene cluster *lngX1, R, S, T, X2, A-J, P*. The gene encoding STh, *estA2* is also, *eatA* and *etpBAC*.

**pAvM_E36_13**

By PlasmidFinder this plasmid was assigned to the less common IncB/O/K/Z plasmid type, and further comparisons revealed that the *repA* gene was 860/873 bp identical to the Z replicon reference (M93064). It is related to a plasmid present in several *Shigella sonnei* genomes as well as a plasmid present in a number of *E. coli* strains that were isolated from faeces (including from that of healthy adults, pCERC10, MF156268) or blood from patients in a hospital.

**Plasmid features:** The plasmid contains a full transfer region and an *oriT* was identified. Several plasmid stability genes are present: *psiAB*, *relE and stbAB*. In addition, the plasmid also has a *pil* locus encoding for a thin pilus. The plasmid harbours the ssRNA *sok* and a putative *hok* gene part of the Hok/Gef protein family.

**AMR:** This plasmid carries a complete Tn*10* containing the *tet(B)* tetracycline resistance genes. IS*10*L of Tn*10* is interrupted by ISSbo1.

**ETEC strain E2980 (L3)**

The ETEC strain E2980 belong to the same lineage, L3, as the CFA/I-expressing strain E36. Strain E2980 express CS7 and LT, specifically LT1. In total three plasmids are present, pAvM_E2980_14-16. Two virulence plasmids and one antibiotic resistance plasmid.

**pAvM_E2980_14**

The virulence plasmid is assigned to IncI1, however one of the marker genes for this Inc group is not present (*sogS*) and the identified marker gene (*trbA*) is not 100% identical to the reference indicating that this may be a new ST allele. This plasmid also contains a *pil* locus with >90 % amino acid similarity with the *pil* operon present in the reference plasmid R64 (AB027308.1). The plasmid is near identical (93-97% coverage and >99% identity) three plasmids all belonging to O144 ETEC strains; strain 90-9276 plasmid unnamed2 (CP024298.1) and strain 90-9280 plasmid unnamed1 (CP024241.1), both collected in Bangladesh 1988, and strain E2264 plasmid unnamed1 (CP023350.1) collected in Bangladesh 18 years later in 2006 (Figure 1d).

**Plasmid features:** A partial *tra* region and several plasmid stability genes, *parA*, *relE* and *stbAB,* are present as well as the ssRNA *sok* is present but the plasmids lack the *hok* and *mok* gene.

**Virulence:** This is a virulence plasmid containing the CS7 locus (*csvA, B, C, E, F, D*) as well as the *eltAB* encoding LT, specifically the variant LT1. The CS7 regulator, CsvR, is also present in this plasmid. In a recent paper it was proposed that the expression of the putative virulence factor *cexE* is translocated with the help from the *aat* locus because the *cexE* gene was located directly downstream of the *aat* operon [12]. However, in this plasmid the *aat* locus is present roughly 17,500 bases upstream of *cexE* adjacent to the regulator *csvR* (Figure 1d).

**pAvm_E2980_15**

This is an IncFII (FII-17, novel allele) resistance plasmid of with a classical IncFII *copA* replication region including an extensive conjugation machinery. The plasmid lacks an *oriT* but the relaxases *traI* is present. The *oriT* may have been lost in the circularisation process as the *traI* relaxases is positioned just at the end of the plasmid. The plasmid is closely related to another plasmid of the above mentioned CS7+ ETEC strain E2264 plasmid unnamed2 (CP023351.1) with 92% query coverage and 99.9% sequence identity. The plasmid is also related to the O25 strain F5505C1 plasmid unnamed1 (81% query coverage and 99.1% identity) which belong to a different ETEC lineage. Similar plasmids are present in other *E. coli* strains.

**Plasmid features:** A complete *tra* region and several plasmid stability genes, *psiAB*, *relE* and *stbAB,* are present. The ssRNA *sok* is present *sok* and a putative *hok* gene part of the Hok/Gef protein family.

**AMR:** This plasmid harbours a 6 kb resistance region, containing a fragment of Tn*5393* containing the *strA* and *strB* resistance genes, *sul2* in a fragment of GIsul2 and $bla_{\mathrm{TEM}}$ derived from Tn*2*. The IR of Tn*5393* bounds one end of the region, while IS*26* is at the other.

**pAvM_E2980_16**

This is a quite small plasmid (48,305bp) in comparison to other plasmids presented here. It was not possible to determine the incompatibility group for the plasmid (pMLST and PlasmidFinder predictions were inconclusive), but is similar to Inc1 and hence called Inc1-like. It is highly similar to a plasmid (CP023352.1) part of the CS7+ ETEC strain E2264 with 94% query coverage and 99.9% sequence identity which has a similar size.

**Plasmid features:** No *oriT* or relaxases was found. The plasmid stability *stbAB* and *vapBC* genes were present in this plasmid. The plasmid also harbours the TA-system *vapBC*.

**Virulence:** This plasmid contains genes encoding two putative ETEC virulence factors, *eatA* and *etpBAC*.

**ETEC strain E1441 (L4)**

E1441 express LT, CS6 and CS21 and is similar to ETEC isolates ATCC 43886/E2539C1 and 2014EL-1346-6 [20]. These isolates were collected in the 70-ties [13] and 2014 (from a CDC collection), respectively, and assigned as O25:H16 which is the O group determined for E1441 *in silico*.

**pAvM_E1441_17**

This is a multi-replicon virulence and resistance plasmid, IncFII (FII-35, novel allele) and IncFIB (FIB-35, novel allele). The plasmid is similar (78% coverage, >99% identity) to a plasmid (CP024258.1) from the ETEC strain F5505-C1 collected 1998 (location unknown).

**Plasmid features:** A 29.3 kb segment of the transfer region has been inverted between two inversely oriented copies of ISCro1b. Despite all of the transfer genes being intact, the functionality of this locus is unknown. A predicted *oriT* was identified. Several plasmid stability genes are present: *stbAB*, *sopAB* as well as the TA-systems *pemI/K.* The ssRNA *sok* is present and a putative *hok* gene part of the Hok/Gef protein family..Furthermore, the *srnAC* plasmid antisense RNA-regulated system [14] is also present.

**Virulence:** The gene cluster *(ΔlngX1*, *lngR, S, T, X2, A-J, P*) encoding the colonisation factor CS21. Notably, several SNPs as well as a 9bp insertion has occurred in the first 31 bases of *ΔlngX1* (in comparison with *lng* locus in pAvM_E925_5). The function of lngX1 is not known, hence whether the mutations and insertion affect the expression of CS21 is difficult to discern.

**AMR:** This plasmid harbours a complex resistance region composed of fragments of transposons. The inverted repeat of a partial Tn*1000* bounds one end. A class 1 integron is located in a partial Tn*21* segment and carries the cassette array *dhfr15* and *aadA1* as well as *sul1* in the 5'-conserved segment. Following IS*186*B and a fragment of Tn*1721* carrying the *tet(A)* tetracycline resistance module, is the mercury resistance module of Tn*21*, bounding the other end of the region [15,16].

**pAvM_E1441_18**

This is an IncFII (FII-106) plasmid that carries several ETEC virulence and putative virulence factors. The same plasmid is present in two O25 ETEC strains, F5504-C1 and ATCC 43886. The plasmids (CP024259.1 and CP024255.1) share >99% identity with 100% coverage (Figure 1e).

**Plasmid features:** Stability genes present are: *parA*, *psiAB* and *stbAB* as well as a *tra* region. The ssRNA *srnC* is present but the *srnB* gene was interrupted by an insertion element (interrupted ISEc12). Another ssRNA, *sok*, was identified.

**Virulence:** The plasmid harbours the gene cluster encoding the CF CS6 (*cssA, B, C, D*) and *eltAB* (LTh). An interrupted *eatA* is present with multiple non-synonymous SNPs and gaps most likely resulting in a non-functional gene. Another putative virulence factor, *cexE*, is located upstream of the *aat* locus (*aatPABCD*) (Figure 1e).

**ETEC strain E1779 (L5)**

The CS5+CS6 ETEC strain E1779 belong to lineage 5 (L5) [1] and is LT+STh positive. CS5 and CS6 positive strains are commonly isolated globally and has been reported to be the most common virulence profile in Bangladesh and India [17,18].

**pAvM_E1779_19**

This is an IncFII (FII-11) virulence plasmid. It is 100% identical to multiple ETEC plasmids with the same virulence profile. The 142 kb plasmid has high similarity and similar plasmid size to several plasmids isolated from ETEC in India, Mali and Gambia as part of the GEMS study and subsequently sequenced [56] including e.g. p504237_142 (CP025863.1), p204446_146 (CP025911.1), and p120899_146 (CP025917.1) as well as the 165 kb plasmid F5176C6 plasmid unnamed1 isolated by CDC in 1997 and the plasmid F5176C6 plasmid unnamed1 isolated in Bangladesh in 2009 [19]. This indicates a high level of conservation over time (Figure 1f).

**Plasmid features:** Several *tra* genes are present, but is not complete. Most likely the transfer genes of other plasmid within the same cell is used for its mobility. The following plasmid

stability genes: *psiAB, repAB,* and *stbAB* are present as well as the TA-systems *ccdAB* and *vapBC*. A ssRNA *sok* is located upstream of a putative gene related to the Hok/Gef family.
**Virulence:** CS5 (*csfABCDEFG*) and CS6 (*cssABCD*) and *estA3/4* (STh). CsfR (often annotated as *csvR,* like the regulator of CS7), a transcription factor known to regulate CS5, is located on the same plasmid but non-adjacent to the *csf* locus. A disrupted *eatA* (split into two CDSs) which is most likely non-functional (Figure 1f)

**pAvM_E1779_20**
The 88,759 bp long plasmid is an IncFII (FII-106) and contain the genes for the ETEC toxin LT. It is 100% identical to unnamed plasmid2 (CP023348.1) from ETEC isolate E2256 isolated in Bangladesh 2006 and *Escherichia coli* strain 204446 plasmid p204446_92 (CP025912.1) collected in Mali 2010. In addition, high similarity to plasmids; p103605_83 (CP025922.1), p120899_76 (CP025918.1), and p204576_83 (CP025909.1), isolated as part of the GEMS study in Gambia 2010, 2012 and in Mali 2010, respectively, was evident. Finally, high similarity to F5176C6 plasmid unnamed2 (CP024669.1) isolated from an O167:H5 ETEC collected by CDC in 1997 was also found confirming earlier findings that LT STh CS5+CS6 isolates can be both O167 and O115 but they share nearly identical plasmids.
**Plasmid features:** An incomplete *tra* regions and the stability genes *psiAB* and *stbAB* are present. The plasmid antisense RNA-regulated system *hok-sok* is present, however the ssRNA *sok* is located in the same orientation as the *hok* and *mok* gene.
**Virulence:** The gene *eltAB* encoding one of the ETEC enterotoxins, LTh, is located on the plasmid.

**pAvM_E1779_21**
This an IncFIIY plasmid that contain plasmid-related stability genes as well as hypothetical genes. No virulence or antibiotic resistance genes are present. It is similar (99% coverage and >99% identity) to plasmid p503440_68 (CP025885.1) of the CS6+CS21 strain 503440 [11].

**pAvM_E1779_22**
This an IncFII (FII-44) plasmid containing a complete transfer region highly similar to the region of R1 [2]. The plasmid matches regions of the plasmid (CP044404.1) identified in the O102 *E. coli* strain NMBU-W10C18 collected from surface water outside of Norway in 2018 and a multi-drug resistant plasmid (CP021536.1) from the *E. coli* strain AR_0119.

**ETEC strain E562 (L6)**
Strain E562 belong to lineage 6 (L6) [1] which encompass CFA/I+STh strains with the novel ON3 genotype [20]. In total five plasmids were identified, three plasmids carrying ETEC virulence and putative virulence genes and one resistance plasmid.

**pAvM_E562_23**
The Inc group analysis was inconclusive but subtyping the plasmid identified a potentially new Inc group related to IncI1 ST288 (for details see Additional file 4). The plasmid is most likely originated from Salmonella strains as several plasmids from different serotypes share ~67% coverage and >96% identity with pAvM_E562_23.

**Plasmid features:** The plasmid contains a full *tra* region as well as the P pili conjugation machinery (*pilI-V*). However, this P-pilis is different from the one found in p26 as it contains pilJ and *pilK* and show no sequence homology. Blastn analysis showed that the pil operon is near identical to the pil operon found on plasmids in Salmonella enterica subsp. enterica. In fact, both the *tra* and *pil* loci are indicated to have originated from Salmonella plasmids. Stability genes *parB, psiAB* and *stbAB* are present. The plasmid also harbours the ssRNA *sok* and a putative hok gene part of the Hok/Gef protein family.
**Virulence:** The genes encoding for the putative ETEC virulence genes EatA and the EtpBAC is present. The genes are located close to each other and there are full and partial IS in the flanking sequence, but there is no strong evidence for how these genes were acquired.

**pAvM_E562_24**

The plasmid harbours two replicons: IncFII (FII-73, novel allele) and IncFIB (FIB-42). The plasmid is similar (94% coverage; >99% identity) to plasmid p504239_155 in the O128 ETEC strain 504239 collected in India 2010 [11], however this plasmid is almost twice the size (155,850bp) compare to pAvM_E562_24 with its 86,655bp.
**Plasmid features:** A partial transfer region is present and the stability genes present are *psiAB* and *stbAB*. The plasmid also harbours the *vapBC* genes, a toxin-antitoxin system involved in plasmid maintenance and the ssRNA *sok* and a putative *hok* gene part of the Hok/Gef protein family.
**Virulence:** The plasmid contains the *lng* locus which encodes CS21.

**pAvM_E562_25**

It belongs to the IncFII (FII-84) replicon and carries multiple virulence genes. It is near identical (100% coverage; >99% identity) to plasmid p504239_101 in the CFA/I STh O128 ETEC strain 504239 part of lineage L6 [21,22] (Figure 1g). Similarity across additional plasmids from CFA/I positive O128 plasmids is also found, however these ETEC strains belonging to a different ETEC lineage (L15) [1,11].
**Plasmid features:** Only *traM* is present, in opposite direction to the predicted *oriT.* The plasmid may be mobilizable with the help of another plasmid. Since the circularisation of the plasmid was not complete there may be genes missing that could be involved in conjugation. Several stability genes present are: *parB, psiAB* (truncated *psiA*), *relE/B* (TA-system), *repAB*. The plasmid also harbours the ssRNA *sok* and a putative *hok* gene part of the Hok/Gef protein family.
**Virulence:** CFA/I locus *cfaABCE* and *estA2* (STh), which are clustered together. The *cfaD* gene is interrupted by a premature stop codon and is identical to *cfaD'* (M55661.1) from a previous study [23]. Whether *cfaD'* is functional or not remains unanswered. Although, the AraC-regulator Rns is also located within the plasmid which might be involved in the regulation of the CFA/I fimbriae. Furthermore, the putative virulence factor *cexE* is located downstream of the *aat* locus, however the *aatD* is located non-adjacent (12kb downstream of *cexE*) to the rest of the *aat* locus (*aatPABC*) (Figure 1g).

**pAvM_E562_26**

This plasmid belongs to the less prevalent IncB/O/K/Z group. More detailed comparisons revealed that it's *repA* gene is 1075/1077 bp identical to that of the K reference plasmid (M93063). Blastn analysis of the pAvM_E672_26 reveals that the same plasmid is found several additional species. It is similar (92% coverage; >98% identity) to p15648-2 from the *Escherichia coli* strain NCCP15648. The NCCP15648 strain was isolated from a patient with diarrhoea and has Shiga-like toxin genes and *eaeA* (intimin) which is involved in adherence. The same type of plasmid is also found in *Shigella flexneri* 2a strain 1508 (pSF150802: CP030917.1) and *Salmonella enterica* subspecies enterica strain H-185(pR805a: MK088173.1).

**Plasmid features:** The plasmid contains a complete *tra* region and is most likely conjugative and may aid in the mobilization of other plasmids. The genes *psiAB* involved in plasmid stability are present as well as the plasmid antisense RNA-regulated system *pndAC* [86].

**Virulence:** A *pil* operon is present, although missing *pilJ* and *pilK*, and it is highly similar to the *pil* operons found in the plasmids mentioned above (99.43-99.52% identity). This *pil* operon was first described in a Shiga-toxigenic *Escherichia coli* and shown to be functional but did not appear to be involved in adherence to human cell lines [24].

**pAvM_E562_27**

This is a resistance plasmid belonging to IncFII (FII-35, novel allele). A plasmid (CP023351.1) in the ETEC strain E2264 is the most closely related plasmid (71% coverage; >99% identity).

**Plasmid features:** The *tra* region present is similar (81% coverage; >96% identity) to the transfer region of R1 [2]. However, *traD* has been interrupted by IS*2*. The stability genes *parB, psiAB* and *stbAB* are present. The plasmid also carries the *hok-sok* plasmid antisense RNA-regulated system is also present.

**AMR:** Several antibiotic and heavy metal resistance genes are clustered together in a region bounded at one end by IS*1R* and IS*26* at the other. A complete Tn*2* containing $bla_{\text{TEM-1b}}$ is inserted in IS*1*R. A partial Tn*1721* containing the *tetA*(A) and *tetR*(A) tetracycline resistance genes and the mercury resistance locus from Tn*21* are also present in this region.

**ETEC strain E1373 (L7)**

The ETEC strain E1373 belong to lineage L7 (L7) encompassing CS6+STp ETEC strains with ST182 [1]. CS6+STp strains with ST-type 182 have previously been identified in the feces of travelers in Guatemala and Mexico [25]. The ETEC strain E1373 also has the same O-serotype (O169) as strains known to have caused multiple outbreaks [26–28].

**pAvM_E1373_28**

This is a multi-replicon plasmid, IncFII (FII84, novel allele) and IncFIB (FIB-56, novel allele). Multiple plasmids from CS6+ ETEC strains are near identical (99% coverage; >99% identity) to pAvM_E173_28. For example, the plasmid pEntYN10 (AP014654.2) of the ETEC strain O169:H41 YN1 isolated in 1991 from a patient with diarrhoea in Japan (Figure 1h). The pAvM_E1373_28 has previously been submitted under a different name (pCss_E1373, LN) but was at the time not manually annotated, except for the virulence genes.

**Plasmid features:** The plasmid does not have a transfer region but may be mobilizable with the help of other plasmids. The stability gene *parB and psiAB* are present. Two copies of the ssRNA *sok* is present.
**Virulence:** The plasmid pAvM_E1373_28 harbours the *cssABCD* operon encoding CS6 as well as the gene (*estA1*) encoding for STp. Interestingly, this plasmid harbours two additional CF-like loci, one CS8-like and a F4-like locus. The similar plasmid, pEntYN10 also harbours the CS6, CS8-like and F4-like locus (Figure 1h).

**pAvM_E1373_29**
This is phage-like IncFIB plasmid shows nucleotide sequence homology to the *E. coli* plasmids AnCo1 (KY515224.1) and AnCo2 (KY515225.1) as well as the *Salmonella typhi* phage-like plasmid pHCM2 (AL513384.1). However, the *roi* (Roi is involved in the lytic growth phase [29]) from pHCM2 has been replaced by *terB*, a tellurite resistance gene.
**Plasmid features:** The plasmid stability genes *psiAB*, however the additional plasmid stability genes *stbAB*, *parM*, *pemK* and *pemI* are missing [30].
**Phage features:** This plasmid does not confer any obvious phenotype upon the bacteria and due to its content should be considered a phage-like plasmid. The plasmid does not encode virulence- or antibiotic resistance determinants, however it contains a full complement of bacteriophage genes and shows a remarkably high synteny to not only Phage-Plasmid pHCM2 but also phage SSU5. The genes shared with pHCM2 and SSU5 encode a cluster of genes similar to lambda-like phage tail fibre proteins, major capsid, collar and other important structural proteins. Furthermore, the Phage-Plasmid encodes a wide variety of putative genes potentially involved in DNA metabolism and replication in bacteria and bacteriophage. Some notable genes identified in pAvM_E1373_29 such as *dnaE*, *ssb*, *rnhA*, *dhfR*, *thyA, nrdAB,* replicative DNA primase-helicase and a DNA primase are all involved in DNA metabolism and replication and likely contribute to a DNA synthesis complex with similarities to those encoded in the T4 bacteriophage genome Other genes identified in pAvM_E1373_29 include a ppGpp synthase/hydrolase that is known to be involved in increased tolerance to antibiotics [31,32].
